# An introgression breakthrough left by an anthropogenic contact between two ascidians

**DOI:** 10.1101/2021.08.05.455260

**Authors:** Alan Le Moan, Charlotte Roby, Christelle Fraisse, Claire Daguin-Thiébaut, Nicolas Bierne, Frédérique Viard

## Abstract

Human-driven translocations of species have diverse evolutionary consequences such as promoting hybridization between previously geographically isolated taxa. This is well-illustrated by the solitary tunicate, *Ciona robusta*, native to the North East Pacific and introduced in the North East Atlantic. It is now co-occurring with its congener *C. intestinalis* in the English Channel, and *C. roulei* in the Mediterranean Sea. Despite their long allopatric divergence, first and second generation crosses showed a high hybridization success between the introduced and native taxa in the laboratory. However, previous genetic studies failed to provide evidence of recent hybridization between *C. robusta* and *C. intestinalis* in the wild. Using SNPs obtained from ddRAD-sequencing of 397 individuals from 26 populations, we further explored the genome-wide population structure of the native *Ciona* taxa. We first confirmed results documented in previous studies, notably i) a chaotic genetic structure at regional scale, and ii) a high genetic similarity between *C. roulei* and *C. intestinalis*, which is calling for further taxonomic investigation. More importantly, and unexpectedly, we also observed a genomic hotspot of long introgressed *C. robusta* tracts into *C. intestinalis* genomes at several locations of their contact zone. Both the genomic architecture of introgression, restricted to a 1.5 Mb region of chromosome 5, and its absence in allopatric populations suggest introgression is recent and occurred after the introduction of the non-indigenous species. Overall, our study shows that anthropogenic hybridization can be effective in promoting introgression breakthroughs between species at a late stage of the speciation continuum.

## Introduction

Anthropogenic hybridizations can arise from human-mediated translocations of species outside their natural distribution range, which promote secondary contact between previously geographically isolated taxa (McFarlane & Pemberton, 2019). Such circumstances enable the initial phase of the hybridization to be studied in a natural context (Grabenstein & Taylor, 2018, Hufbauer et al., 2012, Faust, Halvorsen, Andersen, Knutsen, & André, 2018; Popovic, Matias, Bierne, & Riginos, 2020). Anthropogenic hybridizations thus provide unique opportunities to examine gene flow between species experiencing incomplete reproductive isolation, even at a late stage of the speciation process (Viard, Riginos, & Bierne, 2020).

Species introductions are common and occur at increasing rates in the marine realm (Seebens et al., 2017). Ports and marinas, one component of the increasing marine urbanization, are one point-of-entry of many non-native species (Firth et al., 2016), where they can co-occur with native congeners (e.g., Bouchemousse, Lévêque, Dubois, & Viard, 2016b). Consequently, these habitats are prone to facilitating hybridization between introduced and native species. They provide a suitable system to examine secondary gene flow in case of anthropogenic hybridization. This is illustrated by a recent study from Simon et al. (2020), which documented the presence of a singular lineage, named “dock mussels”, originating from a recent admixture between *Mytilus edulis*, native to the North Atlantic, and *Mytilus galloprovincialis*, native to the Mediterranean Sea. These admixed populations are restricted to port habitats in European waters.

The secondary contact between the solitary tunicates *Ciona robusta* and *Ciona intestinalis* in the English Channel is another case study, but with very different outcomes as compared to *Mytilus* spp. system (Simon et al., 2020). *Ciona robusta* (formerly known as *C. intestinalis* type A; Gissi et al., 2017), native to Asia, has been introduced in the early 2000s in the native range of *C. intestinalis* (formerly known as *C. intestinalis* type B; Gissi et al., 2017), in the English Channel (Bouchemousse, Bishop, & Viard, 2016a). The two species are found in syntopy (Nydam & Harrison, 2010, Bouchemousse et al., 2016b), display similar life-cycles (Bouchemousse, Lévêque, & Viard, 2017), and can be easily crossed in the laboratory (Bouchemousse et al., 2016b; Malfant, Coudret, Le Merdy, & Viard, 2017) despite their high molecular divergence (12% of net synonymous divergence, Roux, Tsagkogeorga, Bierne, & Galtier, 2013, Roux et al., 2016). Successful hybridization nonetheless occurs in one direction only, with *C. intestinalis* as the maternal lineage (Bouchemousse et al., 2016b; Malfant, Darras, & Viard, 2018). Demographic inferences based on few individuals but high number of markers derived from 852 coding sequences (total length of 270kb) suggested the presence of several introgression hotspots between the two species (Roux et al., 2013). However, by using 100 ancestry-informative SNPs, this introgression was later shown to be the outcome of past introgression, far preceding the contemporary secondary contact nowadays observed in the English Channel (Bouchemousse, Liautard-Haag, Bierne, & Viard, 2016c). In Europe, where both species occur in sympatry, Bouchemousse et al. (2016c) have found limited evidence for hybridization (i.e., one F1 hybrid out of 449 individuals), and no sign of contemporary introgression (i.e., no F2s or backcrosses). Thus, despite a high hybridization potential, efficient reproductive barriers seem to prevent hybridization in the wild between the native and non-native species. Although these results are based on low genomic coverage, they suggest that introgression between *C. intestinalis* and *C. robusta* is far less common than in *Mytilus* species. Indeed, admixture was effectively detected in dock mussels using similar genomic coverage (Simon et al., 2020). Nevertheless, high genomic coverage can reveal subtler introgression patterns as exemplified in model systems in speciation (sticklebacks: Ravinet et al., 2018; rock periwinkle: Stankowski et al., 2020; drosophila: Turissini & Matute, 2017), as well as in native-invasive systems (cotton bollworm: Valencia-Montoya et al., 2020; honey bee: Calfee, Agra, Palacio, Ramírez, & Coop, 2020).

In this study, we further explored the genome-wide population structure of the native tunicate *C. intestinalis* in the North Atlantic using a large number of SNPs provided by a ddRAD-sequencing approach. Our study expands on the work conducted by Hudson et al. (2020), which described multiple glacial lineages of *C. instestinalis* within Europe. Here, we aim to evaluate the consequences of anthropogenic hybridization with its congener *C. robusta* that has been introduced in the range of one glacial lineage of *C. intestinalis*, in the English Channel. As a control, and for the sake of comparison, we also examined one population of the native species *Ciona roulei* from the Mediterranean Sea. The species status of *C. roulei* has been repeatedly questioned (Lambert, Lafargue, & Lambert, 1990; Nydam & Harrison, 2010; Malfant et al., 2018), and it might better be described as an isolated population of *C. intestinalis*. Interestingly, *C. roulei* can be found in sympatry with *C. robusta*, also introduced in the Mediterranean Sea. Based on genome-wide SNPs, we 1. recovered the population structure described from previous studies for *C. intestinalis* both at fine and large geographical scales 2. provided genome-wide support for a revision of the taxonomic status of *C. roulei*, and 3. provided the first evidence in favor of recent introgression events from *C. robusta* towards *C. intestinalis* in their contact zone, but not in allopatric populations. However, introgression is restricted to a 1.5 Mb region of chromosome 5. Overall, our study shows that anthropogenic hybridization can be effective in promoting gene flow even between species at a late stage of speciation, but in this case introgression can be restricted to localized breakthroughs in the receiving genome.

## Materials and methods

### Sample collection

We studied 397 individuals of *Ciona* spp., previously sampled across the North Atlantic. The sampling locations are shown in Figures 1B (fine-scale) and 2A (large-scale) with details provided in Table S1 in the Supporting Information file. Most individuals (N=346) are *C. intestinalis* sampled from 22 localities in 2012 by Bouchemousse et al. (2016a), expect for one locality (Jer) that was sampled in 2014 by Hudson, Viard, Roby, & Rius (2016). This sampling includes two localities (REK, Iceland, and NAH, US) where *C. intestinalis* is most likely introduced although its status remains debated (i.e., cryptogenic; See Appendix 1 in Bouchemousse et al., 2016a). The sampling scheme aims at covering the known geographic range of this species, with a focus on the English Channel where the species coexists with its introduced congener *C. robusta*. In addition, 19 specimens of *C. roulei*, native to the Mediterranean Sea, were included, along with 32 individuals from the introduced species *C. robusta*, of which 16 were sampled from the Mediterranean Sea (in sympatry with *C. roulei*) and 16 from the English Channel (in sympatry with *C. intestinalis*).

**Figure 1:**
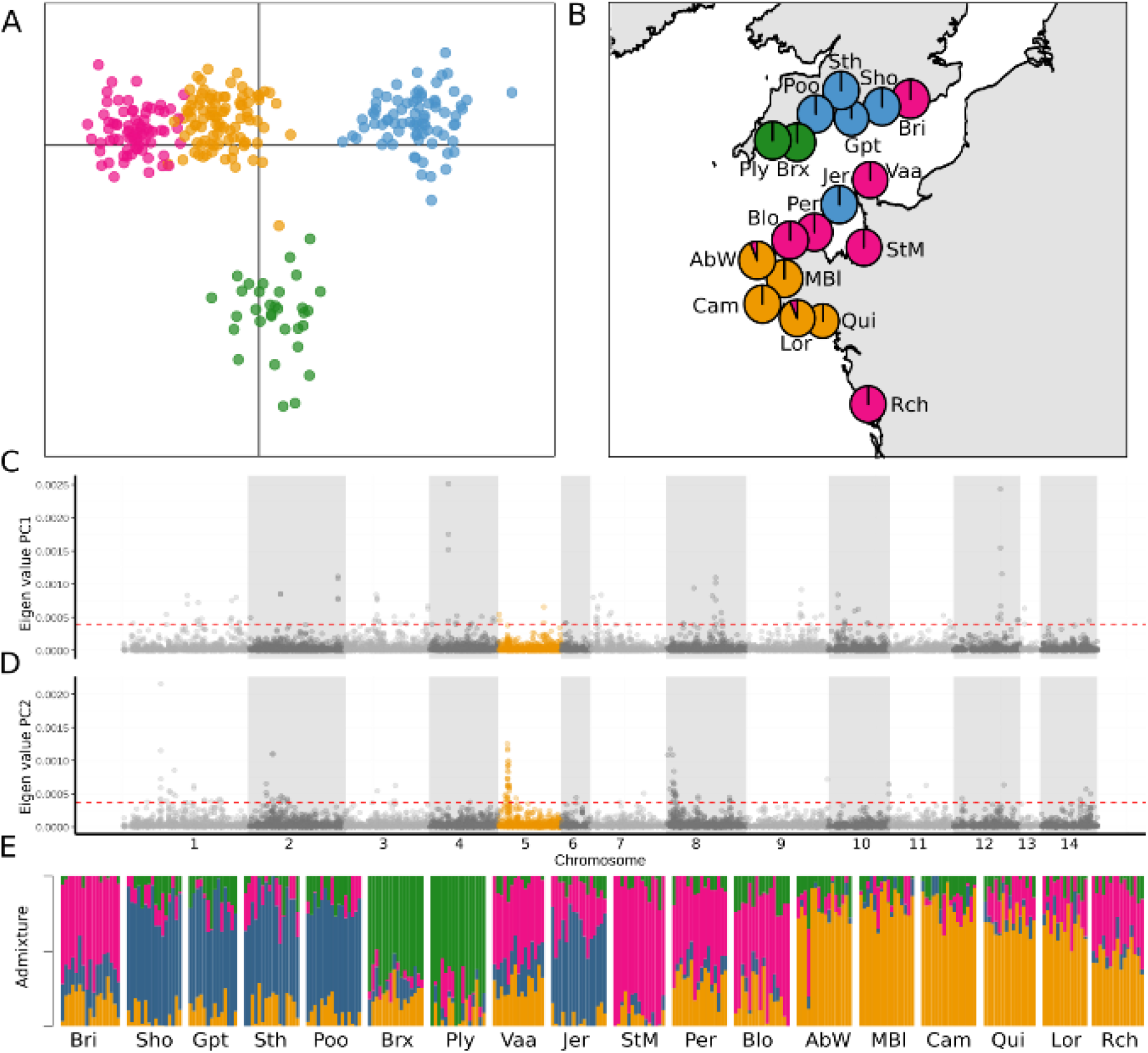
Fine-scale structure analysis of the 18 populations of *C. intestinalis* sampled in Bay of Biscay, Iroise Sea and English Channel. The locality name for each code, and further details about the sampling sites, are provided in Table S1. Based on 280 individuals genotyped at 3,510 unlinked SNPs, four genetic clusters were identified with a DAPC from the find.clusters function, each of them pictured by a different colour in A). Membership to the four clusters is indicated by using the same color scheme in the other plots. B) is showing the proportion of individuals per sampling site assigned to each cluster with the DAPC. The contribution of the 13,603 linked-SNP, mapped on chromosomes, to the first (C) and second (D) axes of the DAPC; the chromosome 5, which is carrying an introgression hotspot, is highlighted in orange; the dashed red lines show the 95% quantiles above which are located the top 5% eigenvalues. E) provides the graphical output of the admixture analyses (*snmf* function of the LEA package; K=4) with the membership of each individual, sorted according to their sampling locality, to each of the four genetic clusters identified with the find.clusters function on the unlinked SNPs.

### DNA extraction and library preparation

For each individual, DNA was extracted using Nucleospin® 96 Tissue Kit according to the manufacturer’s protocol (Macherey-Nagel, Germany). Individual double-digest RAD-seq libraries were constructed with PstI and MseI according to the protocol detailed in Brelsford, Dufresnes, & Perrin (2016), after fluorometric quantification of DNA concentration with PicoGreen (Invitrogen, Carlsbad, CA, USA) and normalization of the extracts. Each individual was labelled with a unique barcode-index combination, with an inline barcode (incorporated in the PstI adaptor) and an Illumina Truseq index (incorporated during the PCR carried out on the ligation products). Size selection was carried out with 1.5% agarose cassettes in a pippin prep (Sage Science) to select fragments between 280 and 600 base pairs. A total of three pooled libraries were sequenced, each containing 184 individuals, with replicates (two individuals per library and two across the three libraries). Each library was sequenced in two lanes of an Illumina HiSeq 2500 v4 high throughput flow cell generating 125 base single-end reads at Eurofins Genomics (Ebersberg, Germany).

### Bioinformatics pipeline

The reads were demultiplexed based on their individual index-barcode with the processRADtags programme of Stacks v2 (Rochette, Rivera-Colón, & Catchen, 2019). Overall, 3% of the reads were dropped due to ambiguous barcodes or low sequencing quality and yield to on average 1.7M reads per sample. The reads were trimmed at 80 base pairs, and mapped on the *C. robusta* genome (KH79 version, Dehal et al., 2002, NCBI assembly GCF_000224145.1) using the default parameters implemented in BWA software (Li & Durbin, 2009). Note that this genome is improperly referenced under the name *C. intestinalis* in several databases because of a recent taxonomic revision (Gissi et al., 2017, and references herein). Between 60 and 67% of the reads were mapped fo*r C. roulei* and *C. intestinalis* individuals, and between 83 and 87% of the reads for *C. robusta* individuals. The aligned RAD data were then processed using the reference mapping pipeline in Stacks v2 set with default parameters (Rochette et al., 2019). Only the SNPs sequenced for at least 80% of the individuals within a locality, present in all the populations and with a maximum heterozygosity of 80% were called using the *population* function. Additional filtering steps were performed using vcftools (Danecek et al., 2011) in order to remove SNPs with a minor allele count of two or showing significant deviations from Hardy-Weinberg equilibrium (P-value threshold of 0.05) in more than 60% of the population with the function *filter_hwe_by_pop*.*pl* implemented in dDocent pipeline (Puritz, Hollenbeck, & Gold, 2014). We additionally removed all polymorphisms private to *C. robusta* populations, as such polymorphisms are neither informative to describe *C. intestinalis* population structure nor to evaluate the extent of introgression across species (which results in shared polymorphisms). At the end of the filtration process, the dataset included 397 individuals (see Table S1 across sampled localities) genotyped at 51,141 SNPs (17,280 with a Minor Allele Frequency (MAF) above 5%) derived from 5,599 RAD-locus with an average depth across all samples of 59 reads per locus, which was then exported into variant call format (VCF). The VCF was then statistically phased using Beagle v5.2 (Browning & Browning, 2007) in order to extract haplotypes and conduct phylogenetic analyses as explained below.

### Population structure analyses

#### Fine-scale population structure (dataset 1)

Because of our interest in identifying introgression in the contact zone, 280 individuals of *C. intestinalis* from 18 sampling sites (70% of the data) were obtained from the English Channel, Iroise Sea and the Bay of Biscay. We first explored the fine-scale population structure of these populations (dataset 1; Table S1; Figure 1B). The dataset 1 was further filtered to keep SNPs with a MAF above 5%, and additionally thinned by keeping one random SNP per over bin of 1kb to take into account physical linkage, using vcftools (Danecek et al., 2011). In total, 13,603 linked SNPs and 3,510 unlinked SNPs were used to study the fine-scale population structure. For the unlinked SNPs, we used the function find.clusters in adegenet (Jombart, Devillard, & Balloux, 2010) to find the best number of clusters (lower BIC value) describing the population structure on the 50 first principal components of a PCA (Figure S1). These clusters were then used as discriminant factors to compute a discriminant analysis of principal components (DAPC) with two discriminant functions. We then used the snmf function of the R package LEA (Frichot & François, 2015), using the number of cluster inferred from the find.clusters function in adegenet to examine the admixture proportions within each locality. We estimated pairwise *F*_ST_ values with 95% confidence interval among sampling sites by bootstrapping (10,000 replications) using the R package StAMPP (Pembleton, Cogan, & Froster, 2013) and significance was tested after accounting for multiple testing with Bonferroni correction. The linked SNPs were used to compute another DAPC using the group inferred by the function find.clusters on the unlinked SNPs, extract the eigenvalue of each individual SNP, and evaluate the genomic distribution of the markers responsible for the population structure.

#### Large-scale population structure (dataset 2)

For the large-scale population structure analyses, and to achieve a more balanced sampling structure, we reduced the number of sampling localities by keeping only one sampling site per genetic cluster inferred with the fine-scale analyses (see above). In total, 129 individuals from nine sampling sites for *C. intestinalis*, 32 individuals from two sampling sites for *C. robusta* and 19 individuals from one sampling site for *C. roulei* were included (dataset 2; Table S1; Figure 2A). This second dataset was filtered to keep SNPs with a MAF above 5%, and further filtered for physical linkage (one random SNP per kb). Overall, 17,138 linked and 3,828 unlinked SNPs were used depending on the analysis. Using the unlinked SNPs, we conducted a PCA analysis from the R package adegenet (Jombart & Ahmed, 2011) and admixture analyses (detailed on Figure S2) from the R package LEA (Frichot & François, 2015). We computed the pairwise *F*_ST_ value with 95% confidence interval from bootstrapping (10,000 replication) using the R package StAMPP (Pembleton et al., 2013), and significance was tested after accounting for multiple testing using Bonferroni correction. A second PCA analysis was performed on the linked SNPs and the eigenvalue of each SNPs were extracted and plotted against their physical position on the reference genome in order to characterize the genomic distribution of the markers responsible for the population structure. Finally, we used the statistically phased VCF file to create a pseudo sequence of all SNPs, and transformed it into a fasta file containing two haplotypes per individual using a custom R script available in the Zenodo archive (see the Data Accessibility Section). From this fasta file, we computed the pairwise genetic distance between each haplotype, and represented a neighbor joining tree based on the GTR substitution model using the R package phangorn (Schliep, 2011).

**Figure 2:**
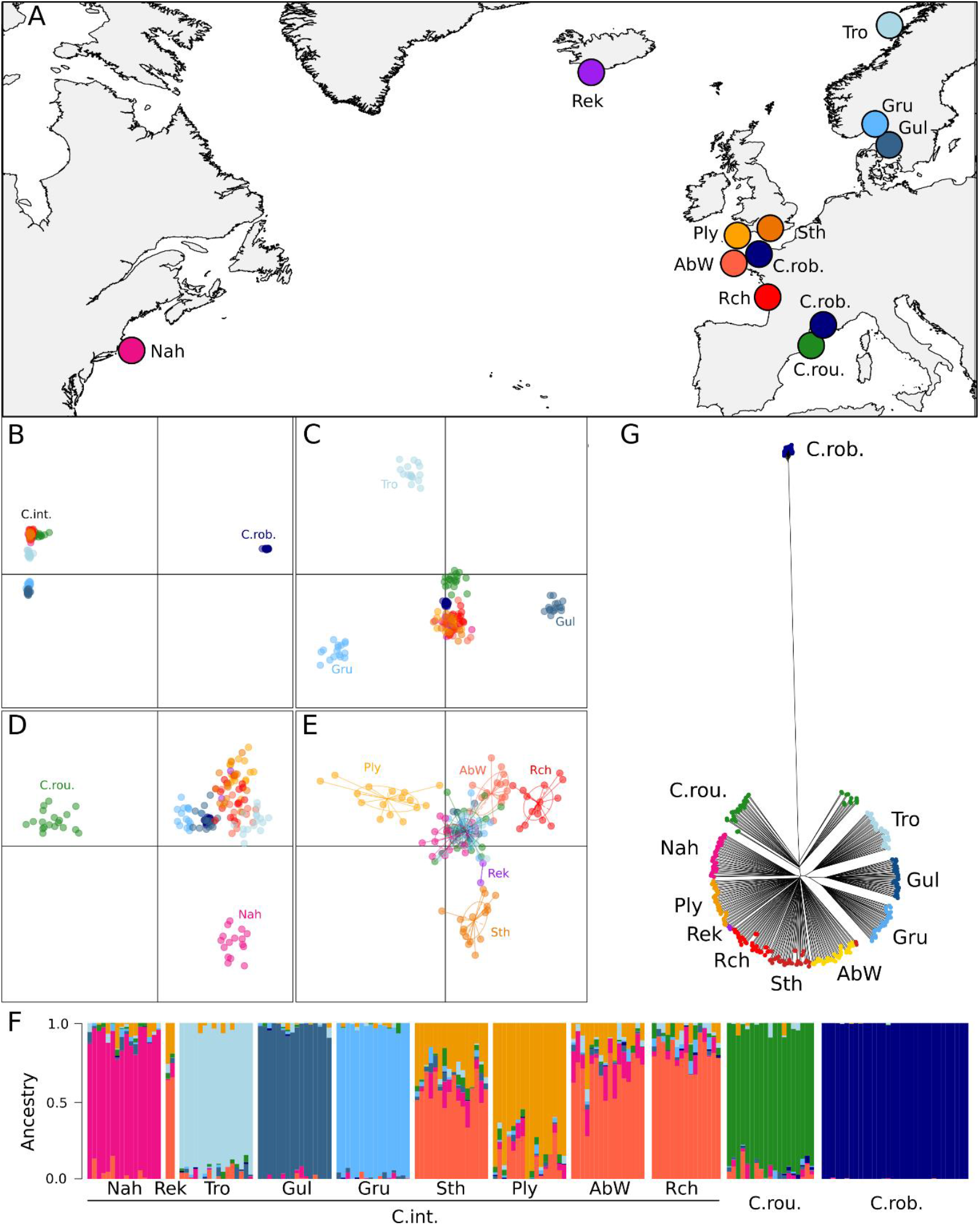
Large-scale population structure analysis of *Ciona intestinalis* (C. int.) populations as compared to *C. roulei* (C.rou.; native to Europe) and *C. robusta* (C. rob., introduced to Europe). The localities examined are shown in A). The locality name, and further details, associated to each code are provided in Table S1. The color code used to picture each sampling locality in A) is used in the other plots. PCA plots, based on 180 individuals genotyped at 3,828 unlinked SNPs, are shown in figures B-E, which are displaying 8 different axes (B: 1 vs. 2; C-E: 3 vs. 4, 5 vs. 6, 7 vs 8), associated to 51.80, 2.56, 1.65, 1.45, 1.29, 0.75, 0.55 and 0.52% of the total inertia, respectively. Figure 2F displays the graphical outcome of the admixture analysis made with the same SNPs using the *snmf* function of the LEA package for a k value of eight cluster (detailed on cluster selection available in Figure S2) for each individual sorted by species and, for *C. intestinalis*, per sampling locality (left part). A neighbor-joining tree of 360 phased haplotype built with the 17,138 linked SNPs is pictured in figure 2G.

#### Investigating introgression between *Ciona intestinalis* and *C. robusta* (dataset 3)

To assess the variability of introgression along the genome, we first used the R package LEA using k=2 to compute the individual ancestry of *C. robusta* and *C. intestinalis* individuals, over the whole genome, and for each chromosome independently. Pairwise *F*_ST_ values between the two sampling localities (pooled) of *C. robusta* and each other sites (*C. intestinalis*) were then computed for each SNP following the method of Weir and Cockerham (1986) using vcftools (Danecek et al., 2011). For each chromosome, we calculated the maximum *F*_*ST*_ value taken over a sliding window of 100kb, which was then smoothed with the R package ggplot2 (Wickham, 2011). Additionally, we created an ancestry informative set of SNPs by extracting the markers differentially fixed (i.e., *F*_ST_ value of one) between *C. robusta* and the Gul population (6,849 SNPs in total). The later population comes from a region and an environment (deep natural habitat) where *C. robusta* have never been reported, thus, is the least likely population to have recently hybridized with *C. robusta*.

To detect potential introgression tracts of *C. robusta* within the genome of *C. intestinalis*, we extracted the *F*_ST_ value of the 6,849 ancestry informative SNPs calculated between *C. robusta* and each 22 *C. intestinalis* populations (Rek excluded due to low sampling size). We then used the Hidden Markov Model (HMM) developed by Hofer, Foll & Excoffier (2012) to infer the position of genomic islands. Briefly, the HMM characterizes and sorts genomic regions according to their level of differentiation, and is generally used to detect island of divergence (e.g. Soria-Carrasco et al., 2014, Shi et al., 2021). Here, the HMM was applied to detect regions of introgression by contrasting regions with high-background differentiation (with an *F*_ST_ value of one) from regions with intermediate differentiation (with *F*_ST_ normally distributed around the 15% lower quantile) and low differentiation (with *F*_ST_ normally distributed around the 5% lower quantile) using a modify version of the R script available from Marques et al. (2016). To avoid any biases by comparing introgressed population from none-introgressed populations, the quantiles were drawn from the distribution of all *F*_ST_ values calculated for the 22 pairwise comparisons. Regions of low differentiation covering more than 4 consecutive SNPs, and regions of intermediate differentiation covering more than 10 consecutive SNPs in a given pair of *C. robusta* and *C. intestinalis* were considered as candidate for introgression. Finally, the genotypes of all the individuals at each diagnostic SNP was visualized using a modified version of mk.image of the R package introgress (Gompert & Buerkle, 2010) available in github and developed as part of Simon et al. (2021). This analysis was performed independently along each chromosome.

## Results

### Fine-scale population structure of Ciona intestinalis in France and UK

Four genetic clusters, which explained 25% of the variance in the DAPC, were identified in the English Channel, Iroise Sea and Bay of Biscay (Figure 1A). The genetic clustering of the localities was consistent with their geographical proximity, except for three of them grouping with geographically distant ones (Figure 1B): 1) the individuals from Jer (Jersey island), geographically close to the northern Brittany populations (deep pink cluster) but clustered with eastern UK sites (blue cluster), 2) Rch (La Rochelle) and Bri (Brighton), which belong to the northern Brittany cluster, despite being geographically closer to Western France (yellow cluster) and eastern UK (blue cluster), respectively. This mosaic structure was also supported by the LEA analysis (Figure 1E), notably for Bri and Jer. However, Rch showed evidence of admixture between nearby localities (Qui and Lor in Western Brittany) and distant ones (Per and Blo in Northern Brittany). Additionally, the two westernmost English populations (Ply and Brx) were genetically differentiated from all the other sampling sites (green cluster in Figure 1B, admixture analysis, Figure 1E).

The SNPs contributing to the population structure observed along the first axis were distributed genome-wide (Figure 1C). Conversely, the SNPs structuring the second axis (and thus differentiating the westernmost English populations) were over-represented at the start of two chromosomes (5 and 8, Figure 1D). Pairwise *F*_ST_ values were overall low (0.001< *F*_ST_ < 0.028) but significant across most sites (Table S2), except among four sites of northern and western Brittany (AbW, MBI, Cam, and Lor), and among four sites of the eastern UK (Sth, Sho, Gpt and Poo). The highest values of *F*_ST_ (0.028) were found among the most distant sites following the coastline, i.e. Rch versus several sampling sites in the UK (Ply, Sho, Poo).

### Large-scale population structure across the Northern Atlantic

The non-indigenous individuals of *C. robusta* are highly divergent from the two species native to European waters, *C. intestinalis* and *C. roulei*, as illustrated on the first axis of the PCA, which explained 51.80% of the inertia (Figure 2B). Populations with less genetic divergence are then distinguished in the following axes: the northern European samples of *C. intestinalis* (Gru, Gul and Tro, in blue) are distinguished by the second axis (2.56%, figure 2B) from the other localities (France, England, Iceland, USA), as well as from the putative sister species *C. roulei*. The following axes (3 to 8) distinguished up to sampling localities of *C. intestinalis* but with much smaller and decreasing inertia, from 1.65% to 0.52% and (Figure 2C-E). *C. roulei* was distinct from *C. intestinalis* only along the axis 5 (1.29%, Figure 2D). The two sampling sites Rek and Nah, in which *C. intestinalis* has an undetermined status (putatively introduced), were very similar to populations from the south of Europe, being distinguished only on axes six and eight (Figure 2D,E).

The SNPs contributing to the major divergence between *C. robusta* and the two native species *C. intestinalis* and *C. roulei* were distributed genome-wide (i.e., Figure S3). However, at the start of the chromosome 5, a reduction of the divergence between *C. robusta* and *C. intestinalis* was observed, as shown by a slight decline of eigenvalues in the PC1 at the start of chromosome 5. The population structure depicted by the PCA was corroborated by the admixture analysis (Figure 2F, Figure S2) showing high support for all the clusters described from the axes one to six of the PCA (Figure 2B-E). All the pairwise *F*_ST_ were significantly different from 0 expect for two comparisons (between the two *C. robusta* sites, and between Rek and Sth, Table S3). The *F*_ST_ values ranged from 0.02 to 0.169 among *C. intestinalis* sampling sites, which is similar to the range (from 0.069 to 0.166) observed between *C. roulei* and any of the *C. intestinalis* populations. The two sites Nah and Rek showed the lowest *F*_ST_ values with the populations from the English Channel (e.g., *F*_*ST*_ Nah vs AbW = 0.033 and *F*_ST_ Rek vs Sth non-significant from 0), and were less differentiated than pairwise comparisons between northern and southern North Atlantic sites. Very large values (i.e., 0.761 to 0.813) were observed between *C. robusta* and *C. intestinalis/C. roulei* (Table S3). The deep divergence of *C. robusta* from the other populations, and that each population clustered in separate groups, is also confirmed by the phylogenetic tree (Figure 2G). C. *roulei* individuals formed a distinct group closely related to *C. intestinalis*.

### Introgression between C. robusta and C. intestinalis

*Ciona robusta* and *C. intestinalis* had consistent high divergence across the genome (Figure S4), with ∼39.64% of the SNPs with a MAF above 5% being diagnostic using Gul population as reference (*F*_ST_=1). No sign of genome-wide admixture was detected between the two species (Figure 3A, top panel), but the *C. roulei* individuals appeared admixed with a *C. robusta* ancestry ranging from 1.96 to 8.77% (Figure 3A, top-panel).

**Figure 3:**
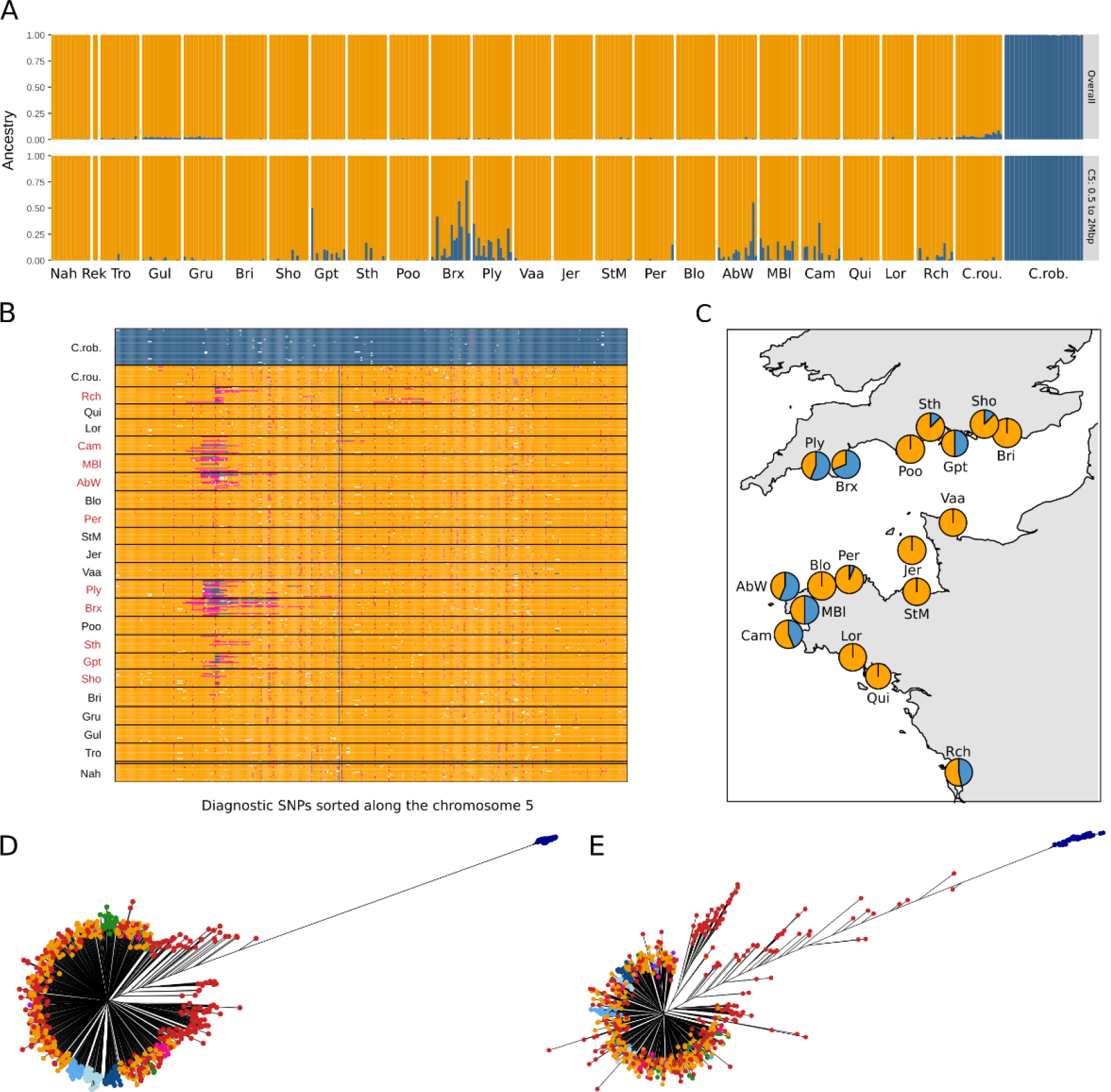
Evidence for introgression, at chromosome 5, of the native species *C. intestinalis* by its introduced congener *C. robusta*, in the English Channel, Iroise Sea and Bay of Biscay. A) Admixture plots computed for K=2 i) on 397 individuals genotyped at 17280 linked-SNPs from the overall dataset (top), and ii) on a subset of 354 SNPs located on the chromosome 5, from 0.5 to 2.0 Mb (bottom). B) Introgress plot showing the genotypes of the 397 individuals at 545 SNPs chosen to be diagnostic between *C. robusta* and *C. intestinalis* along the chromosome 5. Individuals (y-axis) are ordered from top to bottom per species (*C. robusta*: C. rob., *C. roulei*: C. rou., *C. intestinalis*); *C. intestinalis* individuals are sorted per locality (code and location are shown on the map in C)). Population where introgression was detected by the HMM are colored in red. Dark blue boxes indicate homozygote genotype on *Ciona robusta* alleles; yellow, homozygote genotype on *C. intestinalis* alleles; pink, heterozygotes for *C. robusta* and *C. intestinalis* alleles; and white boxes, missing values. C) Proportion of individuals per site displaying a *C. robusta* tract (blue) in chromosome 5. D) and E) Neighbor-joining trees build on 794 phased haplotypes showing the similarities between some *C. intestinalis* haplotypes from admixed localities (red dots) and the haplotypes obtained for *C. robusta* (dark blue dots on the right divergent branch), when using data for chromosome 5 (D) and a zoom from 0.5 to 2.0 Mb of the same chromosome (E); such similarities are not observed for the haplotypes obtained in localities from the contact zone with no introgression (yellow dots), or from other sites located outside the contact zone and from *C. roulei* (colored according to the color code used in Figure 2).

The same results were obtained when each chromosome was analyzed independently (Figure S5), with one noticeable exception found on chromosome 5. On this chromosome, 82 *C. intestinalis* individuals showed a signal of admixture with *C. robusta* (up to 8.62%). The chromosome 5 was also the only chromosome where regions of introgression with a low differentiation between *C. intestinalis* and *C. robusta* at diagnostic SNPs were detected by the HMM, all of which being located between 0.61 and 1.58 Mb (tracts sizes ranging from 64.83kb to 0.49Mb, Table S4). These regions were found in nine populations, and shared an 80 kb fragment located from 0.81 to 0.88 Mb of the chromosome5. Large regions of intermediate differentiation, with tracts sizes ranging from 40.04kb to 1.25Mb, were also found only on chromosome 5, with 90% of them located around the region of low differentiation (from 0.40 and 2.22Mb). In this portion of chromosome 5, the *C. robusta* ancestry reached up to 76.77% (Figure 3A - bottom panel).

The presence of admixed individuals were detected at sites located in the contact zone in the English Channel, Iroise Sea and Bay of Biscay (Figure 3B,C, Table S4). In agreement with the HMM analysis, the chromosome-wide *F*_ST_ values calculated between those populations and *C. robusta* showed a striking decline in *F*_ST_ (Figure 4), with the most extreme drop localized within the 80kb region of low differentiation shared among the most introgressed population, around 0.87Mb of the chromosome 5. The largest decline in *F*_*ST*_ was found in the south western part of the English Channel, in the two populations assigned to the green cluster in the fine-scale analyses (Brx and Ply, Figure 1). Here, *F*_ST_ decreased below 0.5 (Figure 4 – bottom panel), and 29 out of 32 individuals carried at least one *C. robusta* tract (Figure 3A-B-C). The decline in *F*_ST_ was not observed in every sites of the contact zone, being absent in 5 out of 6 of the comparison with populations from the pink cluster identified in the fine-scale analysis (Figure 1), except in Rch (Figure 4). In this latter population, a second *F*_ST_ decline was visible on the same chromosome around 3.1 Mb (Figure 4 – top panel), which is also a region of intermediate differentiation identified by the HMM analysis (Table S4). The admixture signal, and the long tracts of *C. robusta* ancestry were absent from all the sites outside the contact zone, or from other chromosomes (Figure 3A-B, and Table S4), expect in *C. roulei* where few small tracts of introgression (<77kb) were detected on chromosome 7 and 10.

**Figure 4:**
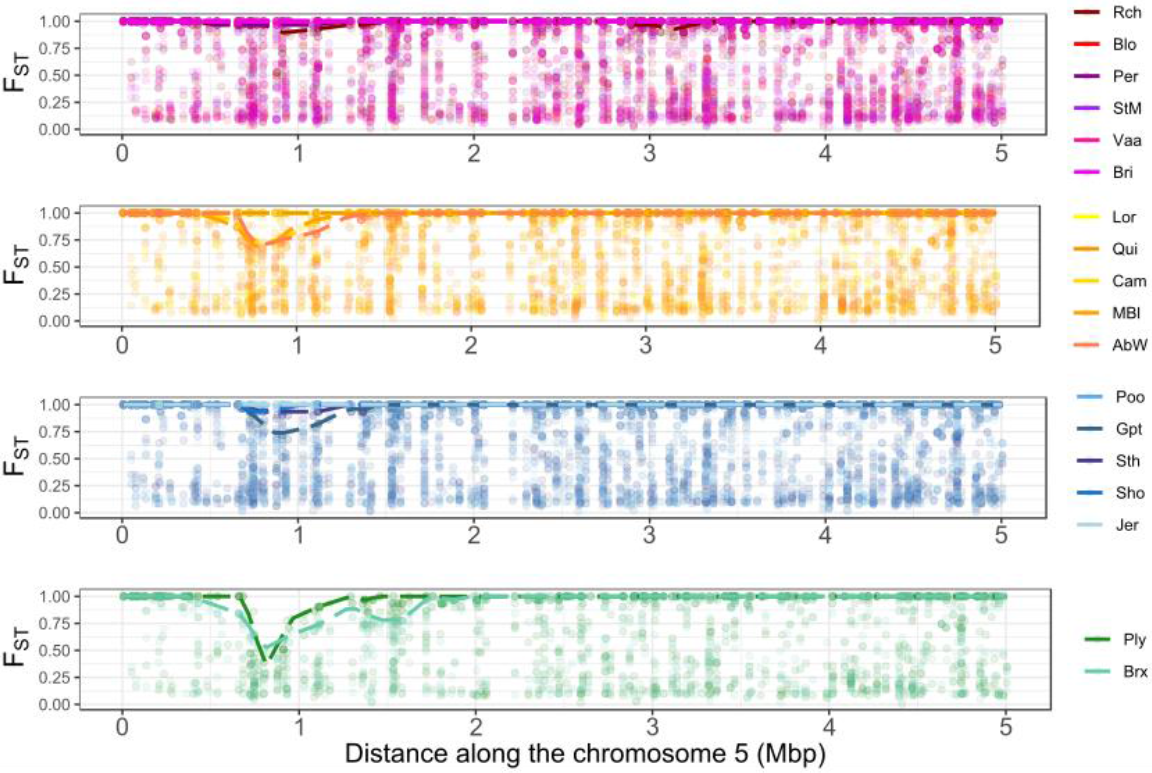
Pairwise *F*_ST_ values along chromosome 5 between *C. robusta* and populations of *C. intestinalis* sampled in the contact zone between the two species, in the English Channel, Iroise Sea and Bay of Biscay. Each graph corresponds to a comparison made with populations from each of the four genetic clusters identified in the fine-scale analyses (clusters are pictured with the same color code as in Figure 1). The populations that belongs to each cluster are listed on the right. Each dot represents the *F*_ST_ value for the 1235 SNP with of MAF of 5% on chromosome 5, and the dashed line show the maximum *F*_ST_ value computed over bins of 100kb.

The neighbor-joining tree built with SNPs from the chromosome 5 showed that some phased haplotypes from admixed sites (red haplotypes in Figure 3D) were genetically closer to *C. robusta* than the haplotypes from non-admixed sites (yellow haplotypes in Figure 3D). This pattern is exacerbated when zooming on the region spanning from 0.5 to 2.0 Mb (Figure 3E). Interestingly, 23 haplotypes (including 10 from 5 homozygous individuals) completely overlapped with *C. robusta* haplotypes when focusing on the portion between 0.7 and 1.2 Mb of chromosome 5 (Figure S6B). In this small region, 19 SNPs were species-diagnostic between Gul and *C. robusta*, and three SNPs were polymorphic in both species (red arrows in Figure S6A). Two of these three SNPs might reflect incomplete lineage sorting or parallel mutation. One was indeed polymorphic in several *C. robusta* and one *C. intestinalis* individual from non-admixed sites (Nah), and the other one was polymorphic in *C. robusta* and in 10 *C. intestinalis* sites, two of which are localized outside the contact zone. Conversely, the last of these three SNPs (position 1020271), was polymorphic in *C. robusta* (maf = 9%), with the minor allele only found in three phased haplotypes of the 23 *C. intestinalis* haplotypes identical to *C. robusta* haplotypes. The three haplotypes carrying the *C. robusta* minor allele at this specific SNP were all collected from the east of UK (one from Ply and two from Gpt). These three haplotypes, different from other *C. intestinalis* haplotypes, all clustered with other *C. robusta* in the phylogenetic tree (black arrow in Figure S6B). Thus, it seems that different *C. robusta* haplotypes have introgressed admixed *C. intestinalis* populations. In addition, the introgression by *C. robusta* was found to be variable in size (Figure 3B) leading to the formation of twin peaks in some pairwise comparison of *C. intestinalis*, as illustrated by pairwise *F*_ST_ values computed between Brx and MBl or AbW (Figure 5C, below the arrows), and also visible when comparing the geographically close sites of Ply and Brx (Figure 5A).

**Figure 5:**
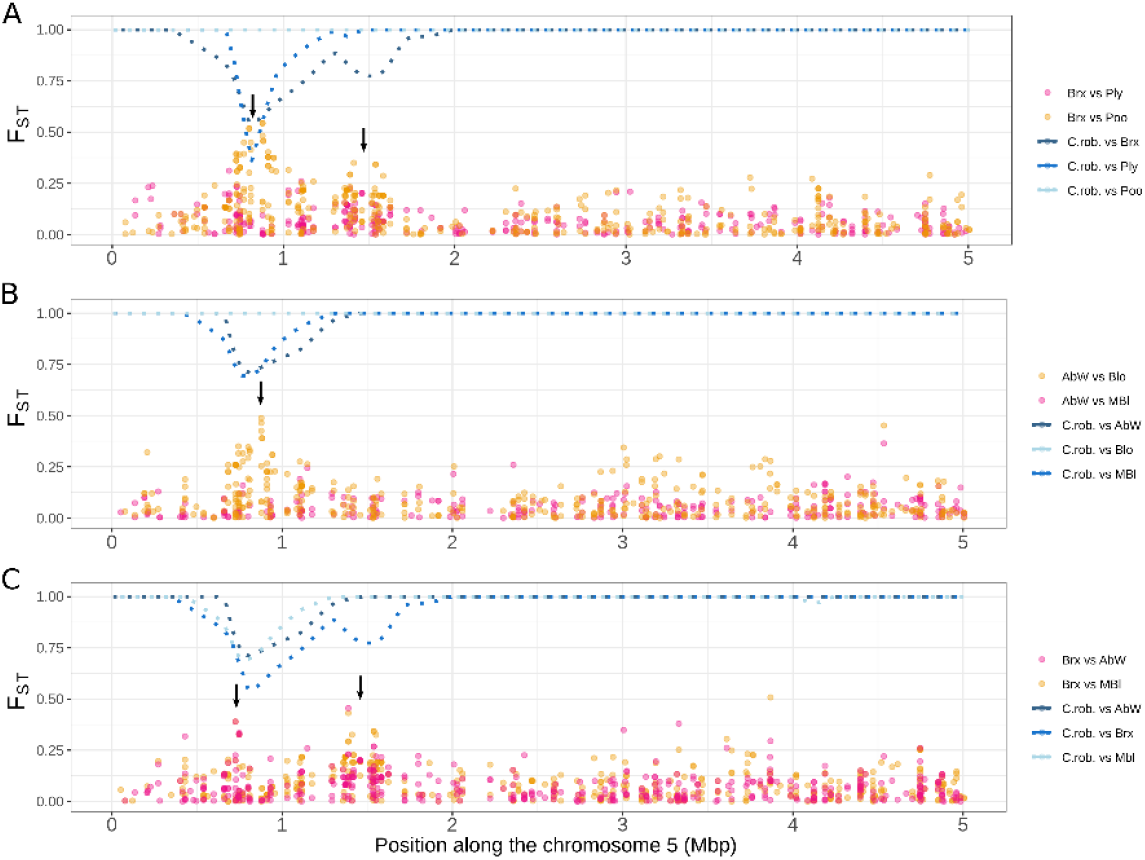
*F*_ST_ values (orange and pink colors) between pairs of *C. intestinalis* populations and maximum value of *F*_ST_ (dashed line in blue colors) between these *C. intestinalis* populations and *C. robusta* over bins of 100kb, computed using 1235 SNPs with MAF of 5% located along the chromosome 5. *F*_ST_ is calculated between pairs of sites either A) geographically close in UK or B) in Brittany (France). In C), *F*_ST_ was calculated between introgressed sites in France vs. UK. Pink color is used to show *F*_ST_ between two introgressed sites and orange color between an introgressed and a non introgressed site.

## Discussion

Using a genome-wide approach based on ddRAD-sequencing, we investigated the fine-scale and large-scale population structure of *Ciona intestinalis* in native (NE Atlantic) and possibly introduced populations (US and Iceland). Our results provide supports to previous studies about the influence of human-mediated transports on population structure at both regional and global scales. Comparisons with its congeners *C. robusta* and *C. roulei* offered new insights about the past and recent history of these three species. Our study finally provided the first evidence of contemporary introgression from *C. robusta* into *C. intestinalis*, in the introduction range of the former. This introgression is not homogeneously distributed in the genome but rather forms a breakthrough located in an introgression hotspot of chromosome 5.

### Chaotic genetic structure and cosmopolitanism: a footprint of human-mediated dispersal

The overall population genetic structure of *C. intestinalis* was in line with the results from previous studies (Bouchemousse et al., 2016a,c; Hudson et al., 2016; Johannesson et al., 2018; Hudson, Johannesson, McQuaid, & Rius, 2020, Johannesson, Le Moan, Perini, & André, 2020). In particular, we confirmed that the populations of *C. intestinalis* are highly differentiated over large geographical scales, which is likely due to the presence of different glacial lineages in Europe (Hudson et al., 2020). At smaller geographical scales, *C. intestinalis* is much less genetically structured, and more importantly shows discrepancies between genetic clustering and geographic distance, leading to a mosaic structure. Such a mosaic structure had been previously reported in the study area, with microsatellite markers, and attributed to human-mediated connectivity among harbors (Hudson et al., 2016). *Ciona intestinalis* is a bentho-pelagic invertebrate but with very short-lived larvae (<24 hours under laboratory conditions). Similar mosaic structures have been documented in introduced species, inhabiting ports, and characterized by low natural dispersal ability, for example the seaweed *Undaria pinnatifida* (Guzinski, Ballenghien, Daguin-Thiébaut, Lévêque, & Viard, 2018).

The individuals sampled in localities for which the native vs. non-native status has been unclear (in the North Western Atlantic and in Iceland) were genetically more similar to the populations sampled in France and England (average *F*_*ST*_ of 0.032) than to other European populations (*F*_*ST*_ up to 0.176). They were also more similar between each other than several comparisons within Europe, suggesting that they are recently derived from somewhere near the English Channel. Such a situation has also been described by Hudson et al. (2020) in a Canadian site localized further north than our study population (Nah). However, the Canadian individuals appeared admixed between Swedish and English Channel lineages, while Nah here appeared as a mostly pure cluster genetically close to the English Channel lineage. Population structure along the western Atlantic Coast is common, even in invasive species such as the green crab (Pringle, Blakeslee, Byers, & Roman, 2011; Jeffery, et al., 2017). The Canadian site is located in Nova Scotia, a major suture zone for marine species living along the Western Atlantic coast, while our sampling site is located further south where populations display usually less admixture (Standley et al., 2018). The differences in admixture pattern observed here and in Hudson et al. (2020) suggest the presence of substantial population structure in the North American introduction range of *C. intestinalis*, which might reflect multiple introduction events from the native range, as often reported in marine introduced species (Viard, David, & Darling, 2016). Additional sampling sites along the Western Atlantic coast would be needed to explore further the population structure of *C. intestinalis*, and reconstruct its introduction history. Our study concurs with previous results by Bouchemousse et al. (2016a) and Hudson et al. (2020), and suggests that *C. intestinalis* can be described as a neo-cosmopolitan species according to the terminology by Darling & Carlton (2018), with a trans-Atlantic distribution due to human introductions, rather than to an eucosmopolitan species with relictual populations that took advantage of new available habitats (ports and marinas).

### A continued and needed appraisal of species status within the *Ciona* genus

The *Ciona* species belongs to a complex genus, as shown by the recent discovery of a new species in the Mediterranean Sea (i.e., *Ciona intermedia*, Mastrototaro et al., 2020), and the recent in-depth taxonomic revision of the previously accepted *C. intestinalis* species. Morphological observations (Brunetti et al., 2015), molecular data (Nydam & Harrison, 2010; Zhan, Macisaac, & Cristescu, 2010; Bouchemousse et al., 2016a,c) and experimental crosses (Lambert et al., 1990; Malfant et al., 2017) indeed showed that two deeply divergent lineages, named type A and type B, were co-existing under the accepted name *C. intestinalis*. This taxonomic revision led to the resurrection of a previously synonymized species, namely *C. robusta*. In line with these previous data, we observed differentially fixed SNPs between *C. robusta* and *C. intestinalis* across most of their genome (average *F*_ST_ > 0.780 for all pairwise comparison). This conclusion also holds for *C. robusta* and *C. roulei*. Our results show that the species status of *C. roulei* is in need of taxonomic revision. We indeed showed a weak genome-wide differentiation when comparing *C. roulei* with *C. intestinalis* populations sampled in the English Channel (*F*_ST_ < 0.075). Interestingly, the differentiation is even weaker than those observed between the southern and northern European populations of *C. intestinalis* studied here (*F*_ST_ up to 0.169). Lambert et al. (1990) and Malfant et al. (2018) performed experimental crosses showing that hybridization between individuals of *C. intestinalis* and *C. roulei* is easy to achieve, with no sign of outbreeding depression. In addition, mitochondrial sequencing data showed that the two taxa display similar haplotypes (Malfant et al., 2018). Our results agree with these previous studies suggesting that *C. roulei* is a divergent lineage of *C. intestinalis*, likely trapped in cold waters of the northern Mediterranean Sea after post-glacial warming, like other cold-adapted marine species (*Platichthys flesus*: Borsa, Blanquer, & Berrebi, 1997; *Sprattus sprattus*: Debes, Zachos, Hanel, 2008; *Sagitta setosa*: Peijnenburg, Fauvelot, Breeuwer, & Menken, 2006).

### Evidence for contemporary introgression of the native species by its introduced congener

Our analyses also provide several novel results. We observed one peak of intra-specific differentiation on chromosome 5 that corresponded to a decline of inter-specific differentiation between sympatric populations of *C. robusta* and *C. intestinalis*. This decline was only found in a subset of the populations located in the contact zone between the two species, pointing toward an introgression from *C. robusta* into *C. intestinalis* populations. The introgression was confirmed by the presence of long *C. robusta* ancestry tracts in some *C. intestinalis* individuals sampled across the English Channel and Iroise Sea, and in the Rch site in the Bay of Biscay. These long introgression tracts were primarily found on chromosome 5 between positions 0.38 to 2.32 Mb (Table S4), which we refer as an introgression hotspot. Other long tracts were found on chromosome 5 outside the main introgression hotspot in a three individuals from Rch and one from Cam. These other tracts close to the introgression hotspot could be either due to independent introgression events or, more likely, to tracts that hitchhiked with the introgression at the hotspot. This introgression restricted to a single hotspot explains why it was missed in previous studies using fewer ancestry-informative SNPs (Bouchemousse et al., 2016b,c). Marker density is thus a key to obtain evidence for very localized introgression (Ravinet et al., 2018, Turissini & Matute, 2017, Stankowski et al., 2020).

*C. roulei* did not show long tracts of *C. robusta* ancestry in chromosome 5, but showed two small tracks (<78kb) on chromosome 7 and 10, and signs of admixture with *C. robusta* were spread across all chromosomes. This admixture pattern and the absence of long haplotypes from *C. robusta* into *C. roulei* could suggest that the introgression between *C. roulei* and *C. robusta* is ancient. Previous studies examining gene flow between *C. intestinalis* and *C. robusta* have interpreted the patterns of allele sharing as a consequence of historical rather than recent introgression (Bouchemousse et al., 2016c). The *C. roulei* samples examined here could thus be another example of historical introgression. Alternatively, such introgression could involve another species closely related to *C. robusta*, but present in the Mediterranean Sea. Out of the 14 species currently accepted in the genus *Ciona* (Word Register of Marine Species; http://marinespecies.org/), three other species not included in this study have been reported in the Mediterranean Sea (*C. intermedia, Ciona* sp. C and *Ciona* sp. D). Based on mitochondrial phylogeny, *Ciona* sp. C appears genetically close to *C. robusta* (Mastrototaro et al., 2020). Thus, carrying out a genome-wide analysis on the *Ciona* taxa found in the Mediterranean Sea is needed to confirm that the signs of admixture of *C. roulei* by *C. robusta* are truly due to introgression with this latter species.

### Spatial and temporal dynamics of the introgression tracts

The recent invasion of *C. robusta* into the English Channel (Nydam & Harrison, 2010), the absence of large introgression tracts outside of the contact zone including in populations outside European Seas (Rek and Nah), the large size of the introgression tracts (>0.5 Mb) and the genetic similarity between introgressed tracts in *C. intestinalis* and *C. robusta* haplotypes (Figure 3), all point toward a recent introgression event (i.e., post-dating the introduction of *C. robusta* in Europe in the late 20^th^ or early 21^st^ centuries). Dating the age of an introgression event is not easy to achieve when using genome reduction DNA sequencing methods, such as ddRAD-sequencing. For instance, Shchur et al. (2020) provided a theoretical framework for interpreting the timing of introgression, based on the distribution of genomic admixture tract lengths, and including positive selection effects. However, the underlying assumption of their method may not hold in our case. For example, the level of genome-wide introgression should be sufficiently high to estimate the baseline neutral introgression rate, while in our case it is around 0.1% on average (Fraisse et al., unpublished results), with little tracts outside of the chrosomosome 5 hotspot. In addition, a single pulse of admixture is assumed in these methods while multiple admixture events have likely happened in *C. intestinalis*, as discussed below.

The localized pattern of long (0.84-1.24 Mb, Table S4), and thus likely young, tracts of introgression in a single region of chromosome 5, and nowhere else in the genome, is difficult to explain without invoking some sort of selection. Although a localized desert of genes associated with low recombination rates can potentially produce this pattern, such regions usually exist at several places of a genome. In addition, 21 genes are located in the 80kb tract shared by the most introgressed populations, from 0.81 to 0.88 Mb of chromosome 5 (listed in Table S5), which does not support the gene desert hypothesis. Moreover, the high variation of *C. robusta* tract length in *C. intestinalis* suggests recombination is operating efficiently on the introgression hotspot. In addition, the introgression is not fixed in any of the study populations. Simple positive selection (adaptive introgression *sensu stricto*) also seems unlikely to explain synchronous incomplete sweeps at many distant locations belonging to different genetic clusters. Alternative selective scenarios therefore need to invoke other kind of selection such as balancing selection, frequency dependence or heterosis. For instance, as the two species studied here are highly divergent (12%, which translates into roughly 2,000 non-synonymous substitutions per Mb), introgression tracts may carry many mutations affecting fitness. Theory predicts an intrinsic benefit of heterozygosity and a cost of admixture (Schneemann, De Sanctis, Roze, Bierne, & Welch, 2020). Introgressed tracts might provide a fitness advantage when heterozygous at the start of the introgression, through heterosis effect, but a negative effect when frequent and homozygote in a *C. intestinalis* background. Furthermore, the heterosis effect could vanish when long tracts are broken down by recombination (Harris & Nielsen, 2016). Thus, the dynamic of the introgression might change over time, with some haplotypes being first positively selected, and then counter selected as soon as recombination broke the introgressed haplotypes into smaller pieces (Leitwein, Duranton, Rougemont, Gagnaire, & Bernatchez, 2020). Examining changes in introgression frequencies and distribution in time would allow to further investigate if selection is truly acting on introgression tracts and continues to drive introgressed tracts to high frequency. Genome sequencing will better delineate the heart of the introgression hotspot and allow to identify candidate genes on the basis of their function.

The length of introgression tracts can be informative about the place where contact happened, as it is inversely correlated to the distance from where the introgression first occurred (Leitwein, Duranton, Rougemont, Gagnaire, & Bernatchez et al., 2020). This effect was documented between the Atlantic and the Mediterranean lineages of European sea bass, *Dicentrarchus labrax*, where the size of the Atlantic introgression tracts into the Mediterranean population is proportionally reduced with the distance from the contact zone between the two lineages, at the Almeria-Oran front (Duranton, Bonhomme, & Gagnaire, 2019). Building on these observations, it is noteworthy that the length of C. *robusta* haplotypes in the genome of *C. intestinalis* individuals was generally larger in western UK (1.24 Mb in Brx, Table S4) than in other populations (e.g., ∼0.5 Mb in Gpt/Sth/MBl), suggesting that introgression occurred there first. Interestingly, introgression was not found restricted to a particular area and shows a chaotic spatial structure similar to what is observed genome wide in our study, and in Hudson et al. (2016). Intra-specific gene flow promoted by human-mediated transports, as previously suggested (Hudson et al., 2016), could explain the chaotic dispersal of introgression tracts. In this context, introgression tracts could be a relevant tool to identify recent human-mediated migration routes (Gagnaire et al., 2015). For instance, while the population Jer seems genetically related to distant populations located in central-eastern UK at the genome-wide level (Figure 2B), this population shares a lack of introgression tracts with StM (Figure 3C), a population in close vicinity, and strongly connected through leisure boating.

Another non mutually exclusive hypothesis might explain both the mosaic structure of introgression and the variable size of introgression tracts: independent introgression events occurring several times in different populations. For instance, the honey bee (*Apis mellifera scutellata*) shows repeated introgressions from the African lineage into the European lineage at one genomic location of the chromosome 1 in two hybrid zones that are localized more than 5,000 kilometers apart (Calfee et al., 2020). Repeated adaptive introgression has been also documented at a gene involved in insecticide resistance in the mosquito *Anopheles* spp. (Weill et al., 2000; Norris et al., 2015) and in cotton bollworm *Helicoverpa* spp. (Valencia-Montoya et al., 2020). Although *Ciona* species are much more divergent than any of the examples presented above, the contact zones between *C. intestinalis* and *C. robusta* are not restricted to a few localities, as they co-occurred in syntopy in several isolated harbors along coastlines of Great-Britain and France, which increases the possibility of introgression. We showed that one SNP distinguished two haplotypes of *C. robusta* found within *C. intestinalis* individuals, suggesting that introgression from *C. robusta* into *C. intestinalis* could have happened at least twice. This SNP was only carried by long introgressed tracts, and could be a footprint of more recent introgression contributing to the twin peak of differentiation observed in the surrounding of the main introgression hotspot between populations that carry introgression tracts of variable size (Figure 5).

The possibility of repeated introgression events does not rule out the hypothesis of intra-specific diffusion of the introgression, facilitated by human activities, and both mechanisms might have jointly contributed to the current distribution of the introgression tracts. Overall, our study shows that anthropogenic hybridization can be effective in promoting gene flow between species even at late stage of speciation, contributing to the population structure described among contemporary populations. Future work should focus on complete genome sequencing, and temporal sampling of the introgressed populations.

## Acknowledgements

The authors are grateful to Sarah Bouchemousse and Sabrina Le Cam for providing samples from Iceland and Norway, and to Alan Brelsford and Pierre-Alexandre Gagnaire for providing advices about ddRAD-sequencing and/or scripts for RAD-Seq analyses. The authors are also most grateful to the Biogenouest Genomics core facility for their technical support, and from the Biogenouest ABIMS Platform for softwares and bioinformatics tools set-up, and access to computing resources. This work benefitted from funding of the French National Research Agency (ANR) with regards the ANR Project HYSEA (no. ANR-12-BSV7-0011).

## Data Accessibility availability

Data will be available upon publication: Trimmed fasta files will be made available in NCBI deposit; the three filtered vcf files used for the fine-scale, large-scale, and introgression analyses, respectively, as well as the R scripts for population genetics and the HMM analyses, and the custom scripts for phylogenetic analyses will be made available on zenodo archives.

## Authors’ contributions

F.V. and N.B. designed the research. F.V. obtained the funding grants. C.D. and C.R. performed the lab-work. C.R. performed the raw-data filtration. A.LM. performed the bio-informatics analyses with advices from N.B. and C.F.. A.LM., N.B., C.F., and F.V. interpreted the data. A.LM. and F.V. wrote the first-draft of the paper, with substantial inputs from N.B. and C.F.. All authors revised and edited the final manuscript.

## Supplemental Information for

**Table S1:**
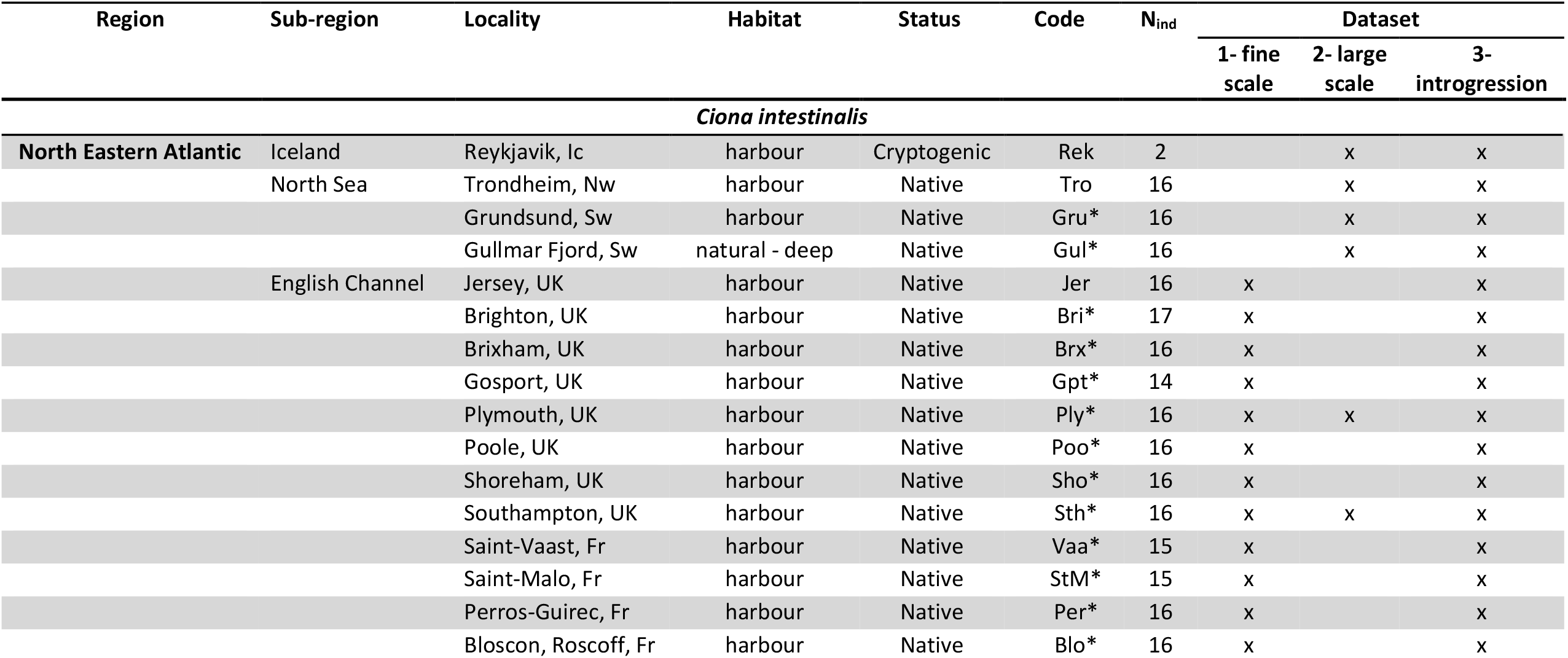
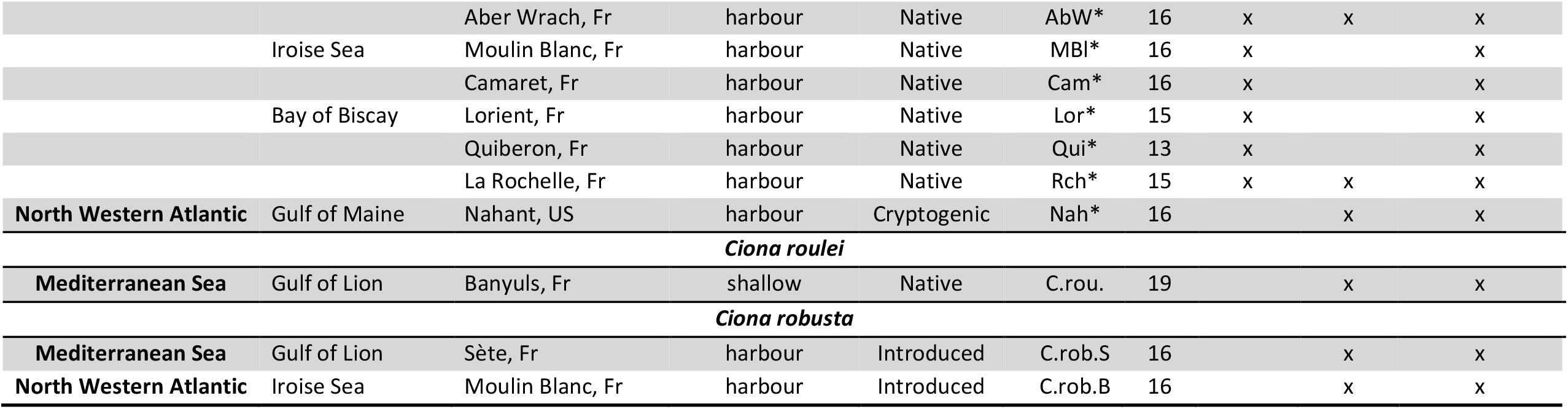
Sampling sites and samples used in the different dataset. For each species, sampling sites (region/sub-region/locality) are indicated with the code associated to each locality, the type of habitat where sampling was made, the species status (native, introduced or cryptogenic), the number of individuals collected, the dataset in which the sampling site was included. “*” in the code column indicates localities which have been examined by mitochondrial sequencing by Bouchemousse et al. (2016a). The Jersey site was collected as part of the study made by Hudson et al. (2016). All the individuals were sampled in marinas from the surface or by diving, except *C. intestinalis* from Gullmar Fjord (Sweden) sampled in natural habitats by diving, and *C. roulei* sampled by diving or trawling in natural habitats but with specimens most often found on artificial substrates, such as tires or watering cans.

| Region | Sub-region | Locality | Habitat | Status | Code | N<sub>ind</sub> | Dataset |  |  |
| --- | --- | --- | --- | --- | --- | --- | --- | --- | --- |
|  |  |  |  |  |  |  | 1- fine scale | 2- large scale | 3- introgression |
| *Ciona intestinalis* |  |  |  |  |  |  |  |  |  |
| North Eastern Atlantic | Iceland | Reykjavik, Ic | harbour | Cryptogenic | Rek | 2 |  | x | x |
|  | North Sea | Trondheim, Nw | harbour | Native | Tro | 16 |  | x | x |
|  |  | Grundsund, Sw | harbour | Native | Gru* | 16 |  | x | x |
|  |  | Gullmar Fjord, Sw | natural - deep | Native | Gul* | 16 |  | x | x |
|  | English Channel | Jersey, UK | harbour | Native | Jer | 16 | x |  | x |
|  |  | Brighton, UK | harbour | Native | Bri* | 17 | x |  | x |
|  |  | Brixham, UK | harbour | Native | Brx* | 16 | x |  | x |
|  |  | Gosport, UK | harbour | Native | Gpt* | 14 | x |  | x |
|  |  | Plymouth, UK | harbour | Native | Ply* | 16 | x | x | x |
|  |  | Poole, UK | harbour | Native | Poo* | 16 | x |  | x |
|  |  | Shoreham, UK | harbour | Native | Sho* | 16 | x |  | x |
|  |  | Southampton, UK | harbour | Native | Sth* | 16 | x | x | x |
|  |  | Saint-Vaast, Fr | harbour | Native | Vaa* | 15 | x |  | x |
|  |  | Saint-Malo, Fr | harbour | Native | StM* | 15 | x |  | x |
|  |  | Perros-Guirec, Fr | harbour | Native | Per* | 16 | x |  | x |
|  |  | Bloscon, Roscoff, Fr | harbour | Native | Blo* | 16 | x |  | x |
|  |  | Aber Wrach, Fr | harbour | Native | AbW* | 16 | x | x | x |
|  | Iroise Sea | Moulin Blanc, Fr | harbour | Native | MBI* | 16 | x |  | x |
|  |  | Camaret, Fr | harbour | Native | Cam* | 16 | x |  | x |
|  | Bay of Biscay | Lorient, Fr | harbour | Native | Lor* | 15 | x |  | x |
|  |  | Quiberon, Fr | harbour | Native | Qui* | 13 | x |  | x |
|  |  | La Rochelle, Fr | harbour | Native | Rch* | 15 | x | x | x |
| **North Western Atlantic** | Gulf of Maine | Nahant, US | harbour | Cryptogenic | Nah* | 16 |  | x | x |
| ***Ciona roulei*** |  |  |  |  |  |  |  |  |  |
| **Mediterranean Sea** | Gulf of Lion | Banyuls, Fr | shallow | Native | C.rou. | 19 |  | x | x |
| ***Ciona robusta*** |  |  |  |  |  |  |  |  |  |
| **Mediterranean Sea** | Gulf of Lion | Sète, Fr | harbour | Introduced | C.rob.S | 16 |  | x | x |
| **North Western Atlantic** | Iroise Sea | Moulin Blanc, Fr | harbour | Introduced | C.rob.B | 16 |  | x | x |

**Table S2:**
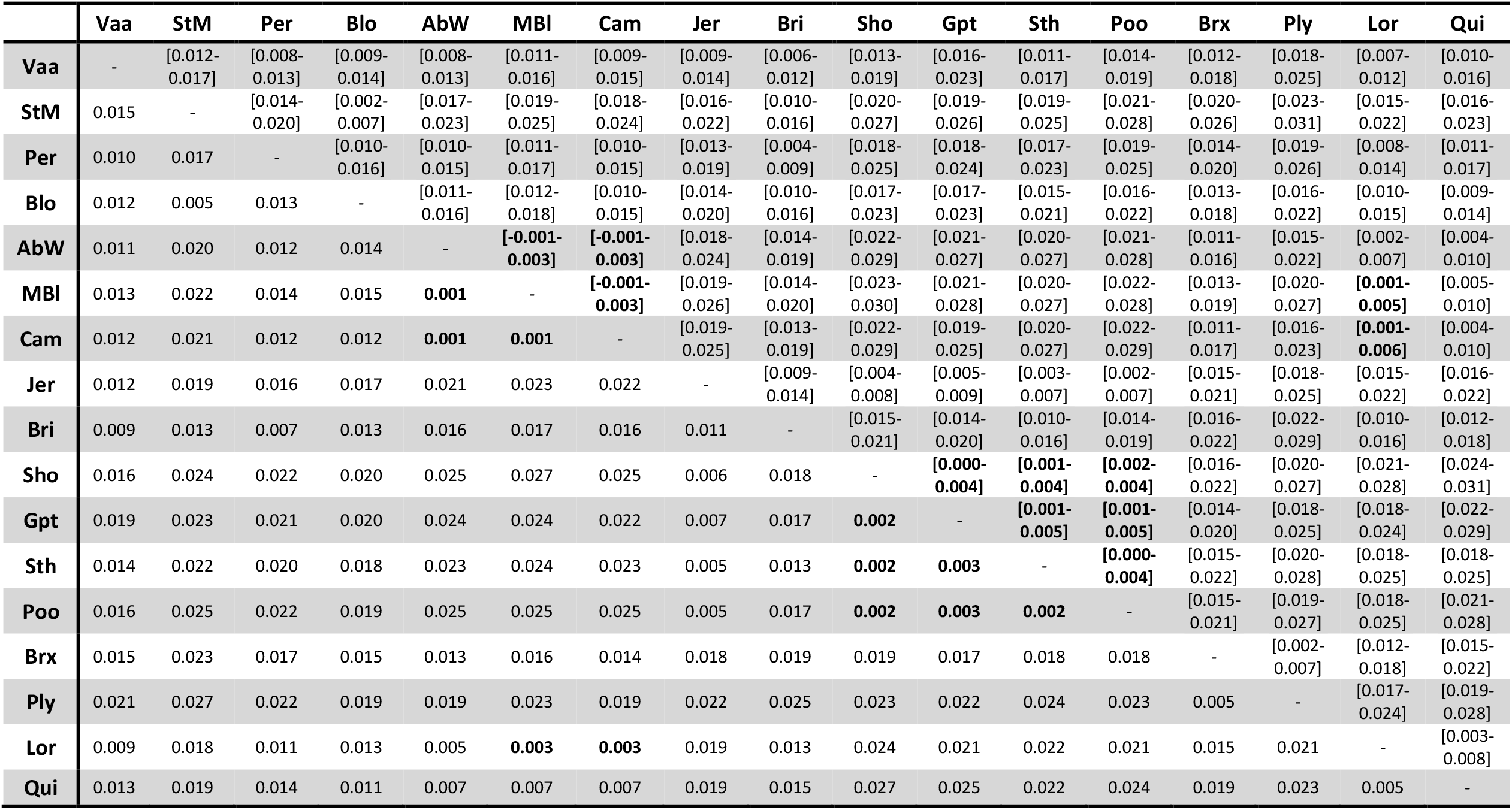
Pairwise *F*_ST_ between populations from the fine scale analyses (bellow the diagonal) with estimates of 95% confidence interval (above the diagonal), non-significant value of *F*_ST_ after Bonferroni correction are highlighted in bold.

**Table S3:** Pairwise *F*_ST_ between populations from the large scale analyses (bellow the diagonal) with estimates of 95% confidence interval (above the diagonal), the non-significant value of *F*_ST_ after Bonferroni correction are highlighted in bold.

|  | Nah | Rek | Tro | Gul | Gru | AbW | Sth | Ply | Rch | C.rou. | C.rob.S. | C.rob.B. |
| --- | --- | --- | --- | --- | --- | --- | --- | --- | --- | --- | --- | --- |
| Nah | - | [0.041-0.066] | [0.102-0.118] | [0.143-0.164] | [0.130-0.150] | [0.029-0.038] | [0.038-0.048] | [0.042-0.053] | [0.038-0.047] | [0.075-0.091] | [0.777-0.796] | [0.777-0.795] |
| Rek | 0.054 | - | [0.104-0.132] | [0.160-0.193] | [0.139-0.172] | [0.012-0.036] | **[0.005-0.027]** | [0.019-0.042] | [0.014-0.039] | [0.051-0.077] | [0.913-0.926] | [0.911-0.924] |
| Tro | 0.110 | 0.118 | - | [0.158-0.179] | [0.145-0.166] | [0.089-0.104] | [0.091-0.106] | [0.095-0.110] | [0.091-0.107] | [0.110-0.128] | [0.790-0.808] | [0.790-0.808] |
| Gul | 0.154 | 0.176 | 0.169 | - | [0.157-0.179] | [0.130-0.151] | [0.133-0.153] | [0.141-0.162] | [0.133-0.154] | [0.155-0.177] | [0.805-0.822] | [0.805-0.822] |
| Gru | 0.140 | 0.156 | 0.155 | 0.168 | - | [0.124-0.145] | [0.121-0.141] | [0.128-0.148] | [0.128-0.149] | [0.146-0.168] | [0.800-0.818] | [0.800-0.817] |
| AbW | 0.033 | 0.024 | 0.097 | 0.141 | 0.134 | - | [0.017-0.024] | [0.016-0.024] | [0.010-0.016] | [0.062-0.077] | [0.774-0.792] | [0.773-0.792] |
| Sth | 0.043 | **0.016** | 0.099 | 0.143 | 0.131 | 0.020 | - | [0.021-0.030] | [0.020-0.027] | [0.063-0.077] | [0.772-0.791] | [0.772-0.790] |
| Ply | 0.047 | 0.030 | 0.102 | 0.151 | 0.138 | 0.020 | 0.026 | - | [0.024-0.034] | [0.067-0.083] | [0.771-0.789] | [0.771-0.789] |
| Rch | 0.043 | 0.026 | 0.099 | 0.144 | 0.138 | 0.013 | 0.024 | 0.029 | - | [0.062-0.076] | [0.781-0.798] | [0.780-0.798] |
| C.rou. | 0.083 | 0.064 | 0.119 | 0.166 | 0.157 | 0.070 | 0.070 | 0.075 | 0.069 | - | [0.751-0.769] | [0.751-0.769] |
| C.rob.S. | 0.787 | 0.919 | 0.800 | 0.814 | 0.809 | 0.783 | 0.782 | 0.781 | 0.790 | 0.760 | - | **[0.003-0.022]** |
| C.rob.B. | 0.787 | 0.918 | 0.799 | 0.814 | 0.809 | 0.783 | 0.781 | 0.780 | 0.790 | 0.760 | **0.012** | - |

**Table S4:**
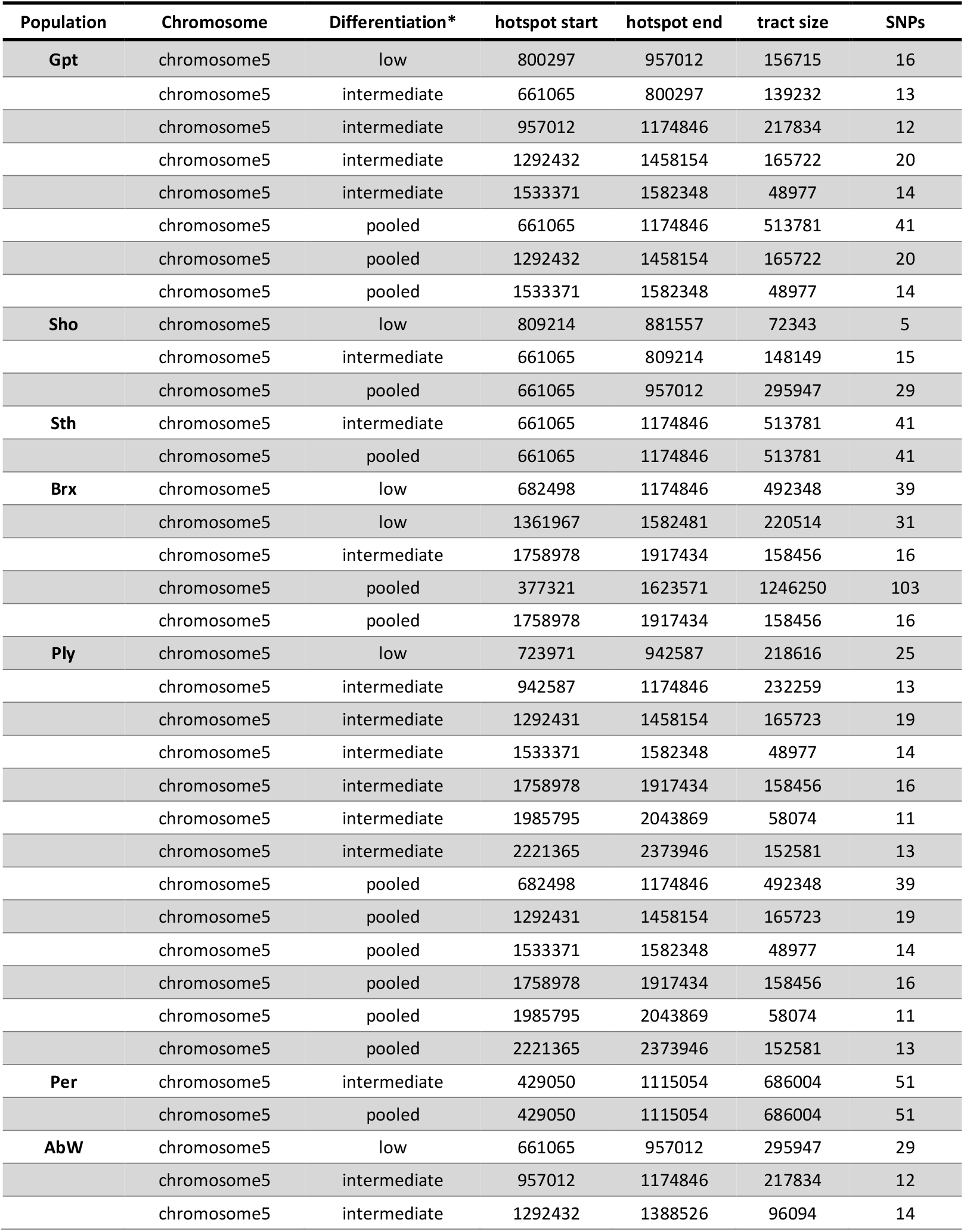
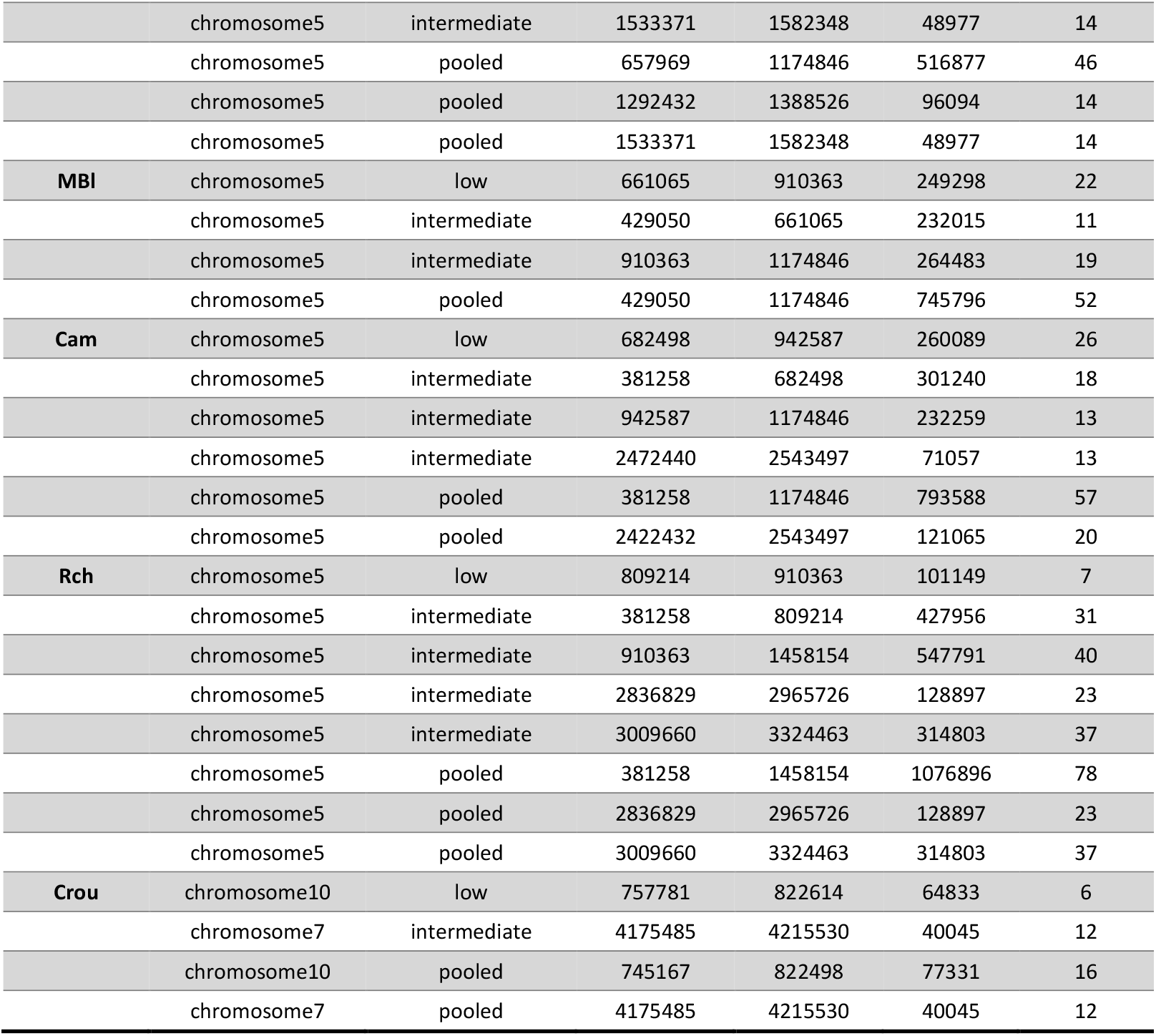
Detection and classification of introgression hotspot from *C*.*robusta* into *C*.*intestinalis* by the Hidden Markov Model (Hufbauer et al., 2012). In order of appearance, the table shows the populations and the chromosomes where introgression hotspots were detected, the differentiation level assigned to the hotspots (“low”= region with at least 4 SNPs with an *F*_ST_ value distributed around the 5% lower quantile, and “intermediate” = regions with at least 10 SNPs with an *F*_ST_ value distributed around the 15% lower quantile, and “pooled” = consecutive regions of “low” and “intermediate” differentiation pooled together), the first base pair of the hotspots, the last base pair of the hotspots, the size of the hotspots and the number of SNPs within hotspots.

**Table S5:**
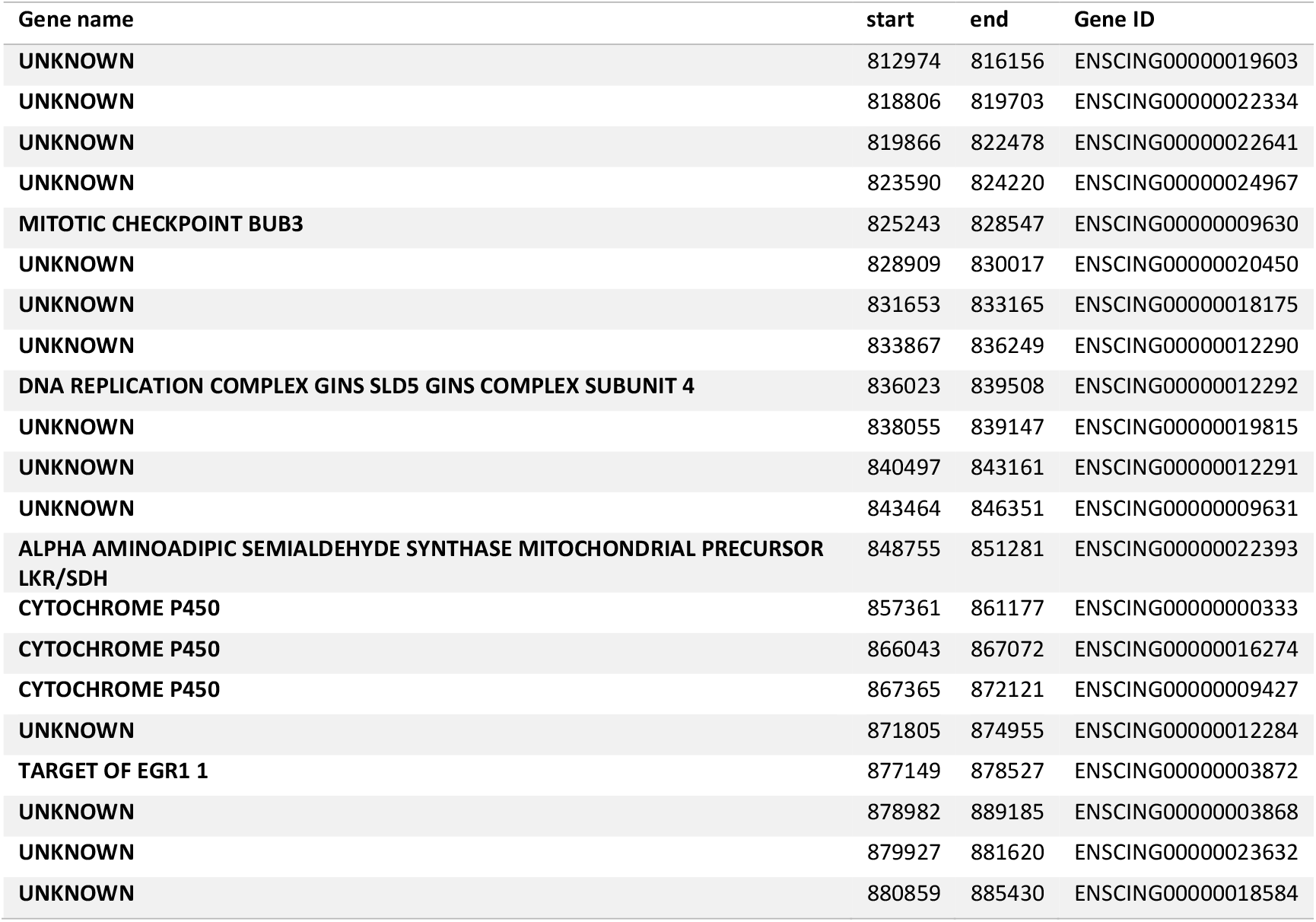
list of the 21 gene localized within the center of the introgression hotspot, i.e. regions constantly found across all the regions of low differentiation in the HMM analyses (see Table S4), from 0.81 to 0.88Mbp of chromosome 5.

**Figure S1:**
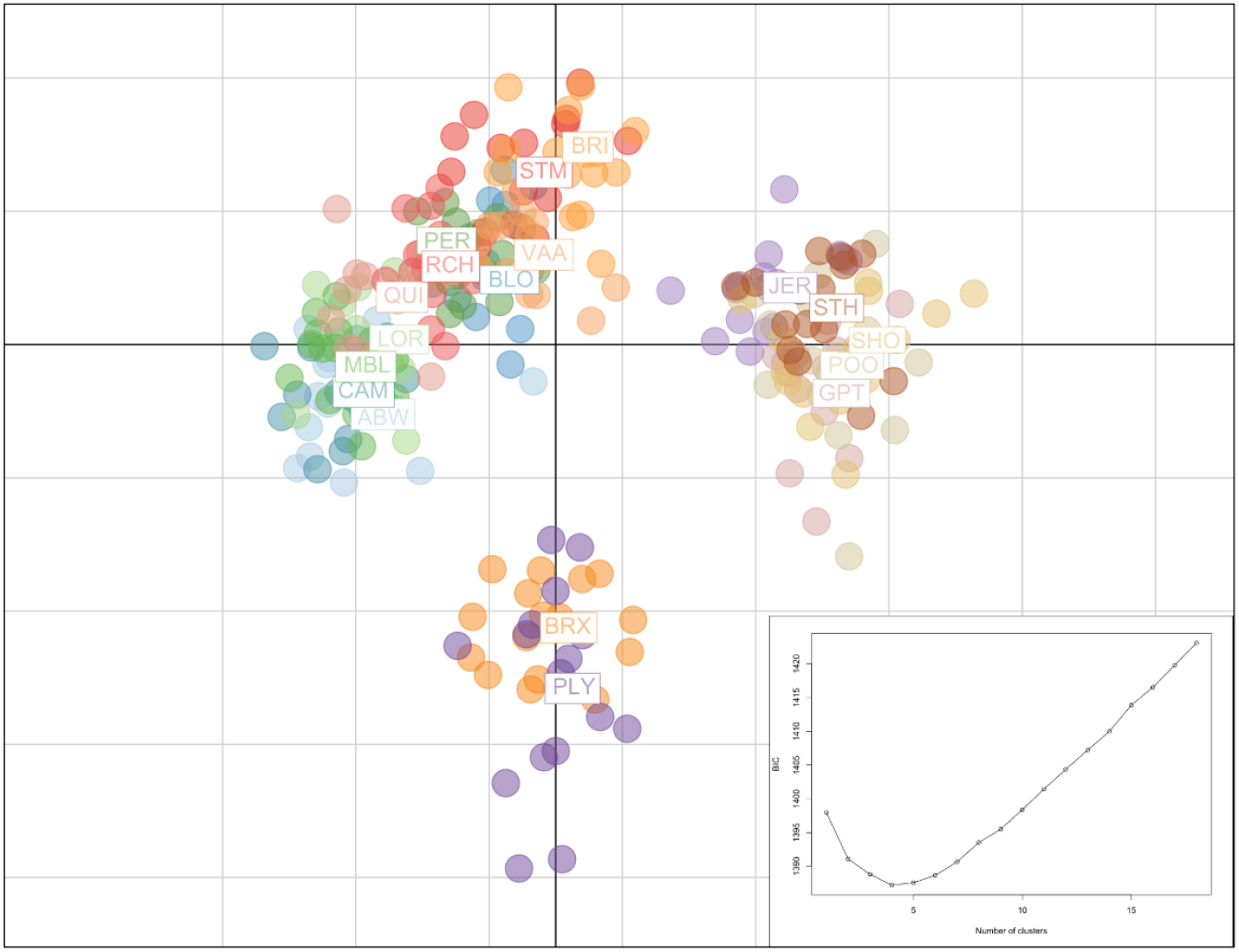
Fine-scale population structure (dataset1) visualized by the two first component of a PCA analyses on the 3510 unlinked SNPs with adegenet (Jombart & Ahmed, 2011). The 18 sampling sites are labeled on the graph according to the code names presented in table S1 and are filled with different colors. PC1 (x-axis) and PC2 (y-axis) showed 1.49% and 0.88% of the total inertia, respectively. Insert shows the value of BIC against the number of clusters assessed by function find.clusters (Jombart et al., 2010). The lowest BIC value is found for a k of four cluster, which was used for the DAPC analyses in the main text in Figure 1.

**Figure S2:**
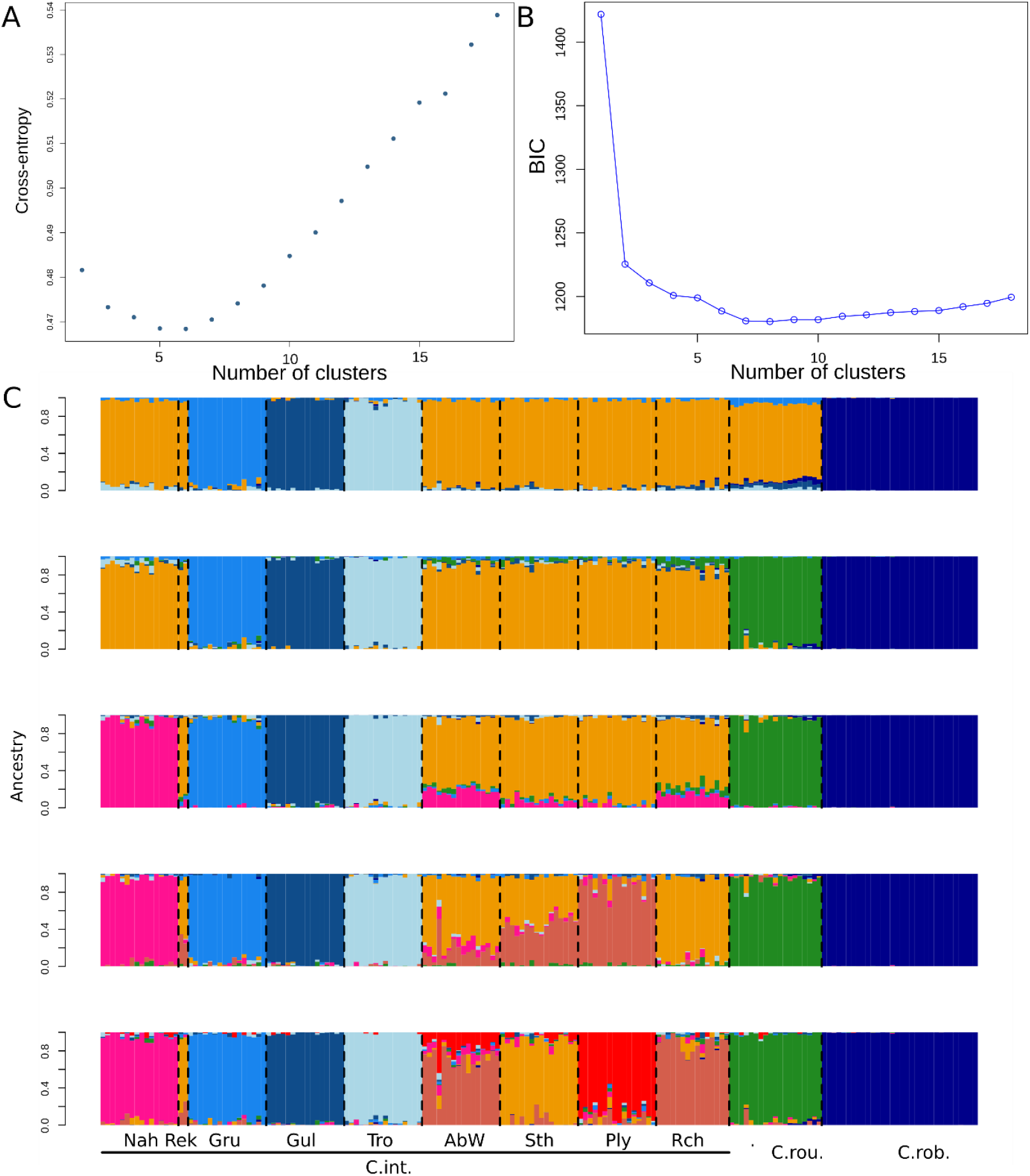
Population clustering for the large scale population structure analyses (dataset2) with A. the cross-entropy validation performed by LEA (lowest value found for a k=6 clusters), and B. BIC variation against the number of clusters by the find.clusters function of adegenet package on the 50 first PC (lowest value found for a k=8) and C. the ancestry proportions inferred by the snmf function of the LEA package for a K value ranging from 5 (top panel) to 9 (bottom panel). The ancestries inferred for K=8 are showed in Figure2 in the main.

**Figure S3:**
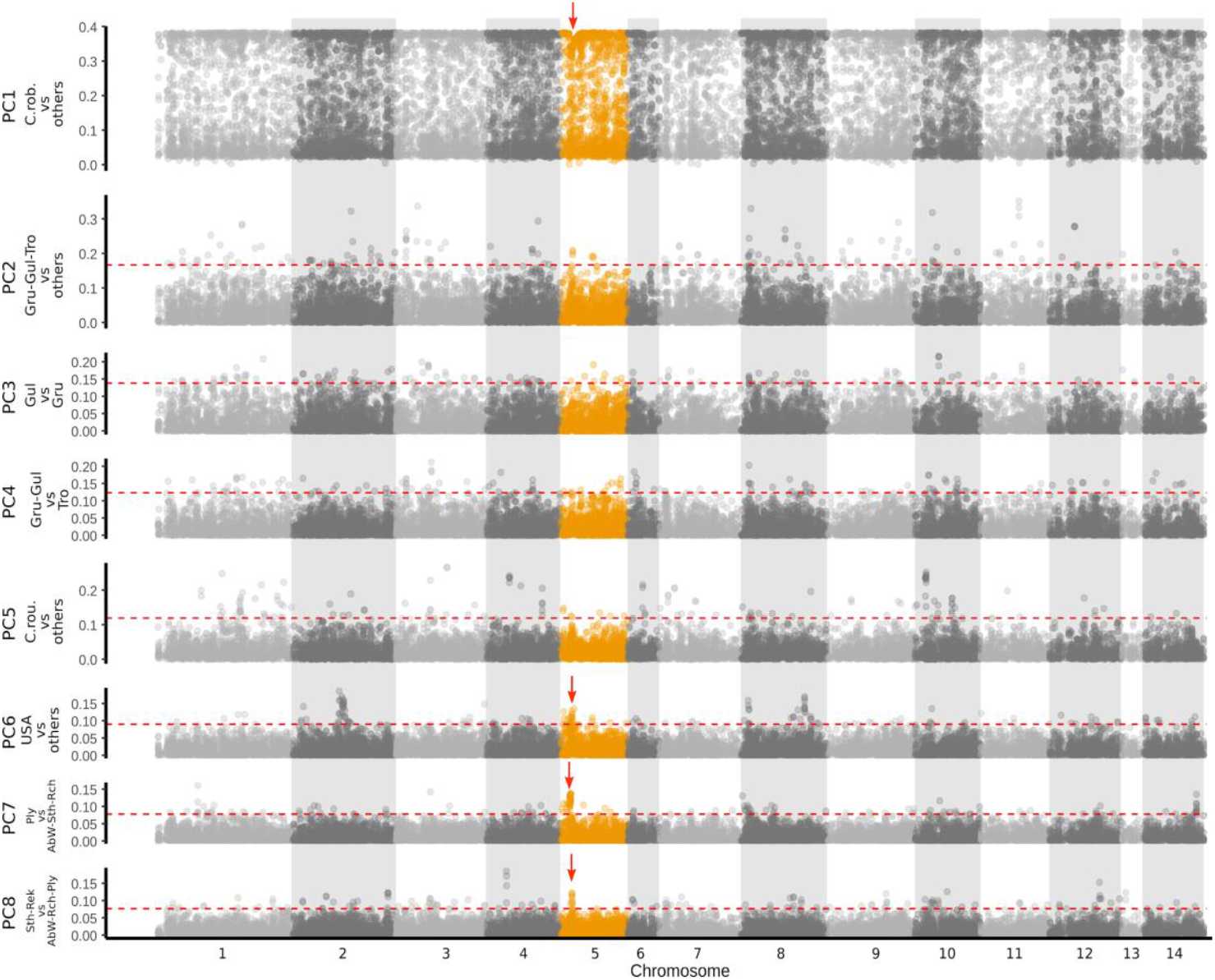
Contribution of each SNP to each axis of the PCA in the large scale analysis shown in Figure 2 (Main text). The red dotted lines represent the 95% quantile above which the top 5% eigen values are located. The chromosome 5 is highlighted in orange, where the red arrows point toward a slight decline of eigen values for the PC1, indicative of a reduction in the genome-wide divergence between *C. robusta* and *C. intestinalis*. The location of this decline is coinciding with a peak of eigen value among the populations of the English Channel and US (red arrows on plots for PCA6-8).

**Figure S4:**
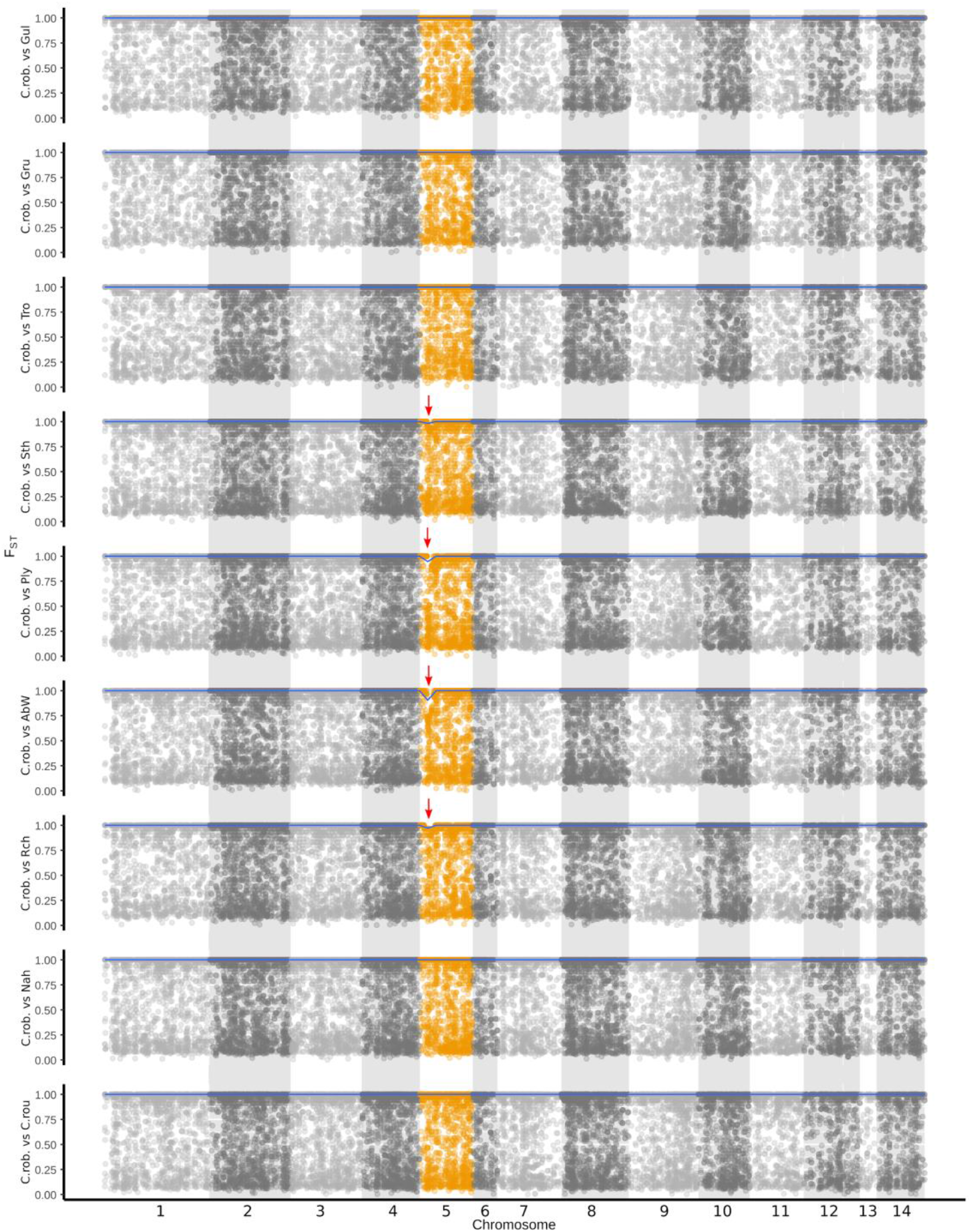
Manhattan *F*_ST_ plot for different pairwise comparison between *C. robusta* and populations of *C. intestinalis* sampled over the large geographical scale. The blue lines represent the maximum value of *F*_ST_ over bins of 100kb. The chromosome 5 is highlighted in orange, where the red arrows point toward a slight decline of *F*_ST_ between *C. robusta* and *C. intestinalis* sampled from the contact zone (Sth, Ply, ABW and Rch).

**Figure S5:**
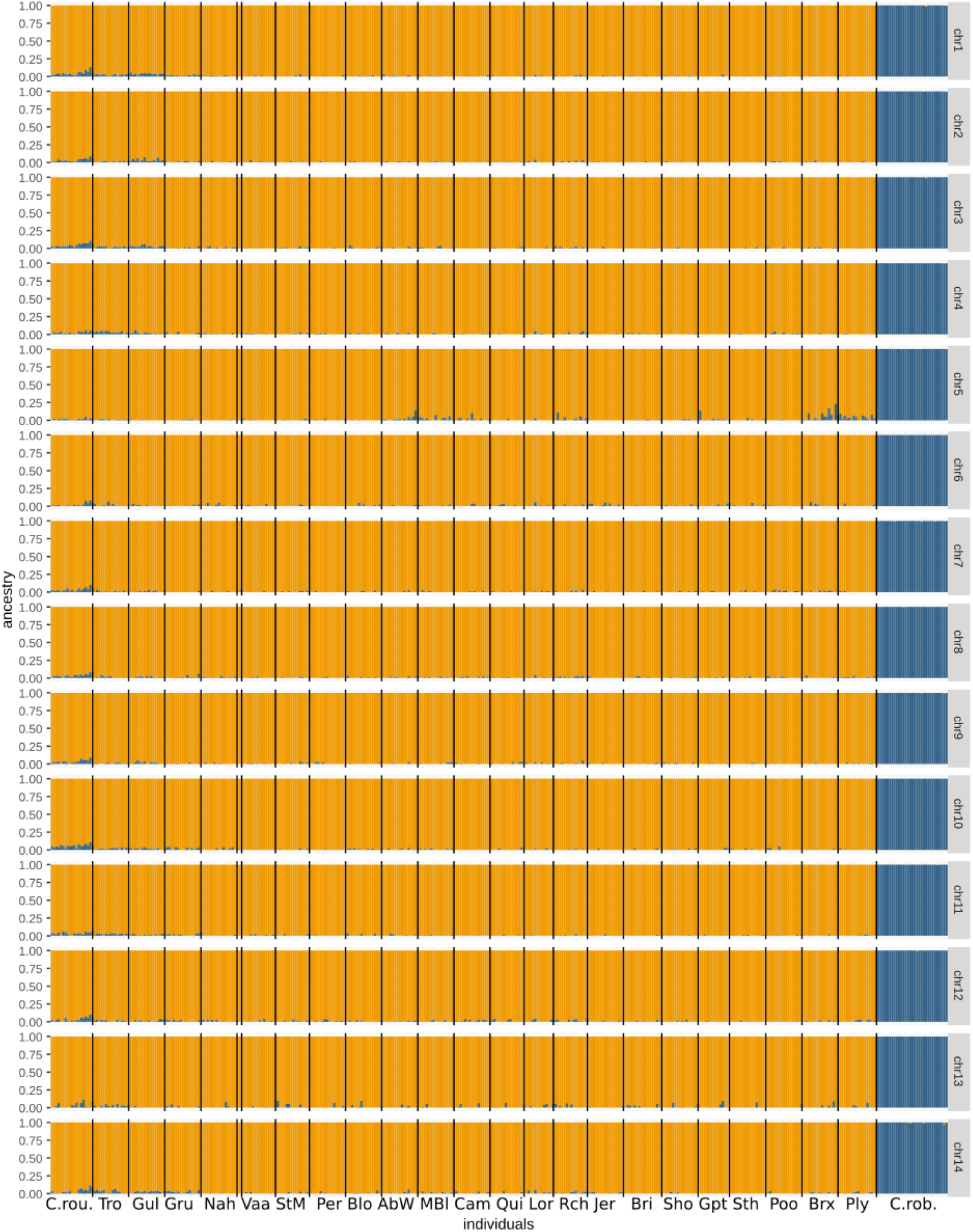
Structure plot per chromosome for k=2 (one plot per chromosome). The vertical lines delimitate individuals collected from different sampling sites. Signal admixture, albeit low, is detected in every chromosome between *C. roulei* and *C. robusta*. However, sign of introgression between *C. intestinalis* and *C. robusta* is detectable only chromosome 5 in 82 individuals sampled from Per, Blo, AbW, MBl, Cam, Rch, Gpt, Sth, Brx and Ply.

**Figure S6:**
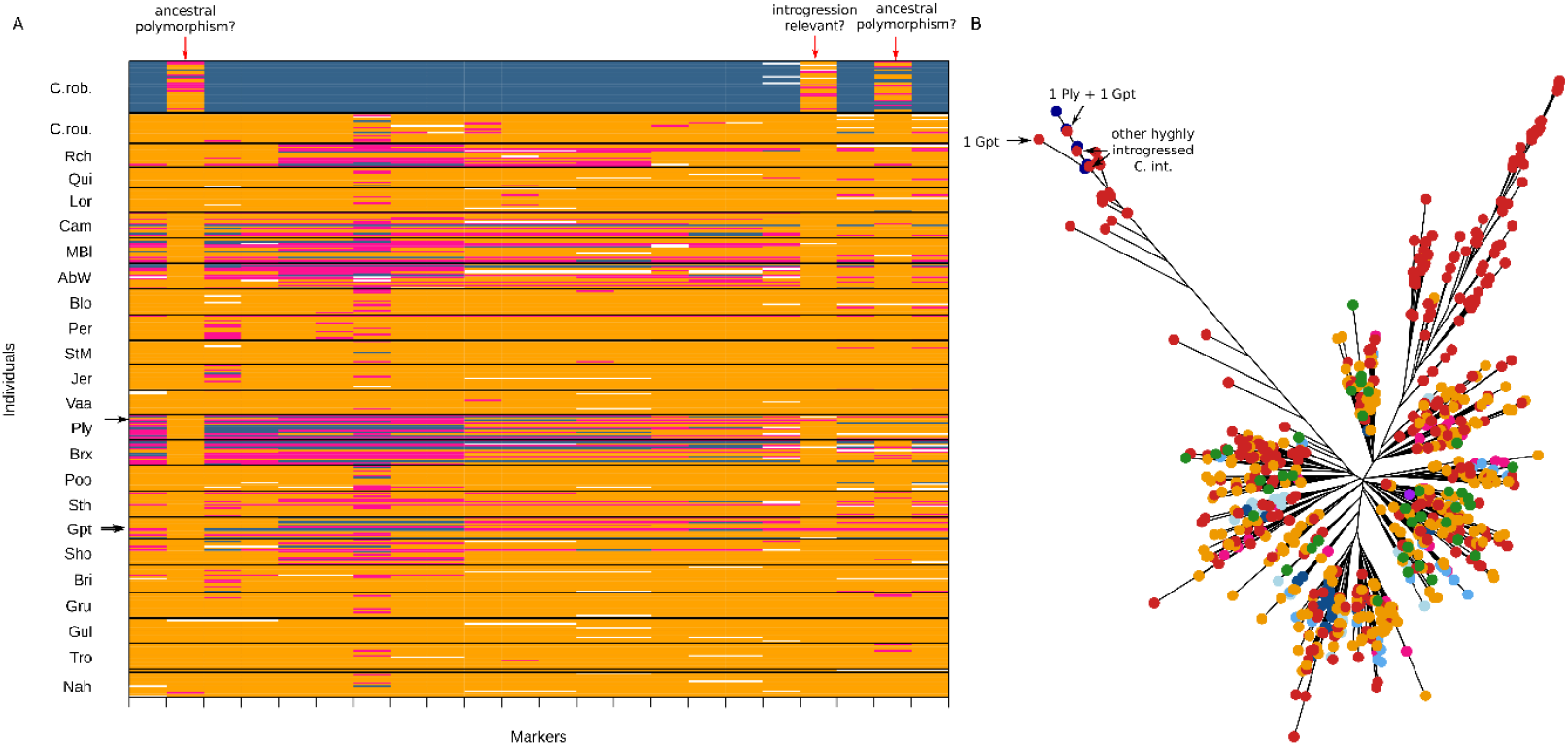
Zoom between 0.7 and 1.2 Mbp of the chromosome 5, with A) Introgress plot for 22 diagnostic and polymorphic SNPs between *C. robusta* and *C. intestinalis* from Gul (locality from deep waters of Sweden). Polymorphism private to *C. intestinalis* and/or *C. roulei* was removed from the analyses. Markers (x-axis) are ordered following physical position on chromosomes. Individuals (y-axis) are ordered per population. Dark blue boxes indicate homozygote genotype on *Ciona robusta* alleles; yellow, homozygote genotype on *C. intestinalis* alleles; pink, heterozygotes for *C. robusta* and *C. intestinalis* alleles; and white boxes, missing values. The plot shows shared polymorphism between *C. robusta* and all *C. intestinalis* individuals (below the red arrow), and B) neighbor-joining tree of all phased polymorphism data where black arrows distinguish different *C. intestinalis* haplotypes (in red) segregating among *C. robusta* haplotypes.

